# Low oxygen enhances trophoblast column growth by potentiating the extravillous lineage and promoting LOX activity

**DOI:** 10.1101/669796

**Authors:** Jenna Treissman, Victor Yuan, Jennet Baltayeva, Hoa T. Le, Barbara Castellana, Wendy P. Robinson, Alexander G. Beristain

## Abstract

Early placental development and the establishment of the invasive trophoblast lineage take place within a low oxygen environment. However, conflicting and inconsistent findings have obscured the role of oxygen in regulating invasive trophoblast differentiation. In this study, the effect of hypoxic, normoxic, and atmospheric oxygen on invasive extravillous pathway progression was examined using a human placental explant model. Here, we show that exposure to low oxygen enhances extravillous column outgrowth and promotes the expression of genes that align with extravillous trophoblast (EVT) lineage commitment. By contrast, super-physiological atmospheric levels of oxygen promote trophoblast proliferation while simultaneously stalling EVT progression. Low oxygen-induced EVT differentiation coincided with elevated transcriptomic levels of lysyl oxidase (*LOX*) in trophoblast anchoring columns, where functional experiments established a role for LOX activity in promoting EVT column outgrowth. The findings of this work support a role for low oxygen in potentiating the differentiation of trophoblasts along the extravillous pathway. Additionally, these findings generate insight into new molecular processes controlled by oxygen during early placental development.

**Summary Statement:** Low oxygen promotes extravillous trophoblast differentiation

## INTRODUCTION

In mammalian development, the placenta forms the mechanical and physiological link between maternal and fetal circulations. In rodents and humans that have invasive haemochorial placentae, nutrient and oxygen transfer between mother and fetus is achieved through extensive uterine infiltration by placenta-derived cells of epithelial lineage called trophoblasts (Pijnenborg et al., 2011; Velicky et al., 2016). In humans, trophoblast differentiation into invasive cell subtypes, called extravillous trophoblast (EVT), is essential for optimal placental function (Tilburgs et al., 2015; Velicky et al., 2016). Molecular processes governing invasive EVT differentiation and specific EVT functions like uterine artery remodeling and immuno-modulation of maternal leukocytes are strictly controlled (Pollheimer et al., 2018; Wallace et al., 2012). Defects in trophoblast differentiation along the EVT pathway associate with impaired placental function and certain aberrant conditions of pregnancy that directly impact fetal and maternal health (Avagliano et al., 2012).

In early pregnancy, anchoring villi of the placental basal plate initiate cellular differentiation events leading to the formation of EVT. At specific villi-uterine attachment points, proliferative EVT progenitors establish multi-layered cellular structures called anchoring columns (Haider et al., 2016; Pollheimer et al., 2018). Trophoblasts residing within proximal regions of anchoring columns, termed proximal column trophoblasts (PCT), are highly proliferative and show evidence of initial molecular characteristics that are hallmarks of EVT (Haider et al., 2018; Turco et al., 2018). At distal regions within anchoring columns, column trophoblasts lose their proliferative phenotype and express many markers akin to invasive EVT, such as HLA-G, α5 and β1 integrins, NOTCH2, and ERBB2 (Fock et al., 2015a; Fock et al., 2015b; Haider et al., 2016; Kabir-Salmani et al., 2004; Zhou et al., 1997). The transition of PCT into distal column trophoblasts (DCT) represents a significant developmental step towards the formation of uterine-invading and tissue-remodeling EVT.

Anchoring columns of the placenta initially develop in the absence of maternal blood, and subsequently, within a relatively low oxygen environment (∼20 mmHg) (Jauniaux et al., 2000; Jauniaux et al., 1999; Rodesch et al., 1992). By comparison, the partial pressure of arterial blood oxygen is approximately 100 mmHg, while the partial pressure of oxygen within the placental bed by 14 weeks’ gestation is estimated to be between 40-60 mmHg (Jauniaux et al., 2001). The role of oxygen in controlling anchoring column formation and EVT differentiation has been the focus of many studies (Caniggia et al., 2000; Genbacev et al., 1997; James et al., 2006; Lash et al., 2006; Wakeland et al., 2017). Unfortunately, contradictory and inconsistent findings have obscured the role of oxygen tension in controlling aspects of trophoblast biology, where low oxygen both promotes (Caniggia et al., 2000; Genbacev et al., 1997) and restrains (James et al., 2006; Lash et al., 2006) column outgrowth, and potentiates (Wakeland et al., 2017) and inhibits (Caniggia et al., 2000; Genbacev et al., 1997) EVT differentiation. These reported differences on the effect of oxygen on anchoring column outgrowth and EVT differentiation are likely attributed to differences in model platforms and methods used to isolate, characterize, and culture trophoblasts. Nonetheless, the role of oxygen in controlling anchoring column formation and column trophoblast differentiation has yet to be fully examined.

In this study, we examine how differing levels of oxygen affect first trimester placental column outgrowth. Using placental villous tissue explants that recapitulate many of the morphological and molecular events central to anchoring column formation and EVT differentiation *in vivo*, we show that low levels of oxygen potentiate column outgrowth. We demonstrate that low oxygen drives hypoxia-related gene programs and processes central to cell-extracellular matrix interaction, while exposure to high oxygen promotes/maintains a strong proliferative phenotype. Moreover, we provide evidence that supports a role for low levels of oxygen in promoting the differentiation of column trophoblasts along the EVT pathway. We further identify *LOX* as a critical gene up-regulated in response to low oxygen that supports column outgrowth, and provide important insight into novel molecular programs impacted by oxygen that align with trophoblast differentiation along the EVT pathway.

## RESULTS

### Hypoxia potentiates trophoblast column outgrowth

To test the effect of exposure to differing levels of oxygen on trophoblast column establishment and outgrowth, we utilized a Matrigel-imbedded placental explant model that closely reproduces developmental processes of trophoblast column cell expansion and differentiation along the EVT pathway (Bilban et al., 2009; Newby et al., 2005). Early column formation during human placental development is characterized by the expansion of mitotically active Ki67^+^/HLA-G^lo^ proximal column trophoblasts (PCT) into HLA-G^hi^ non-proliferating distal column trophoblasts (DCT) and invasive EVT (Figure 1A, 1B). To this end, Matrigel-imbedded placental explants recapitulate anchoring column cell organization and EVT-lineage commitment, establishing the explant system as an appropriate experimental tool to study the cellular and molecular underpinnings regulating human trophoblast column formation and outgrowth (Figure 1C).

**Figure 1.**
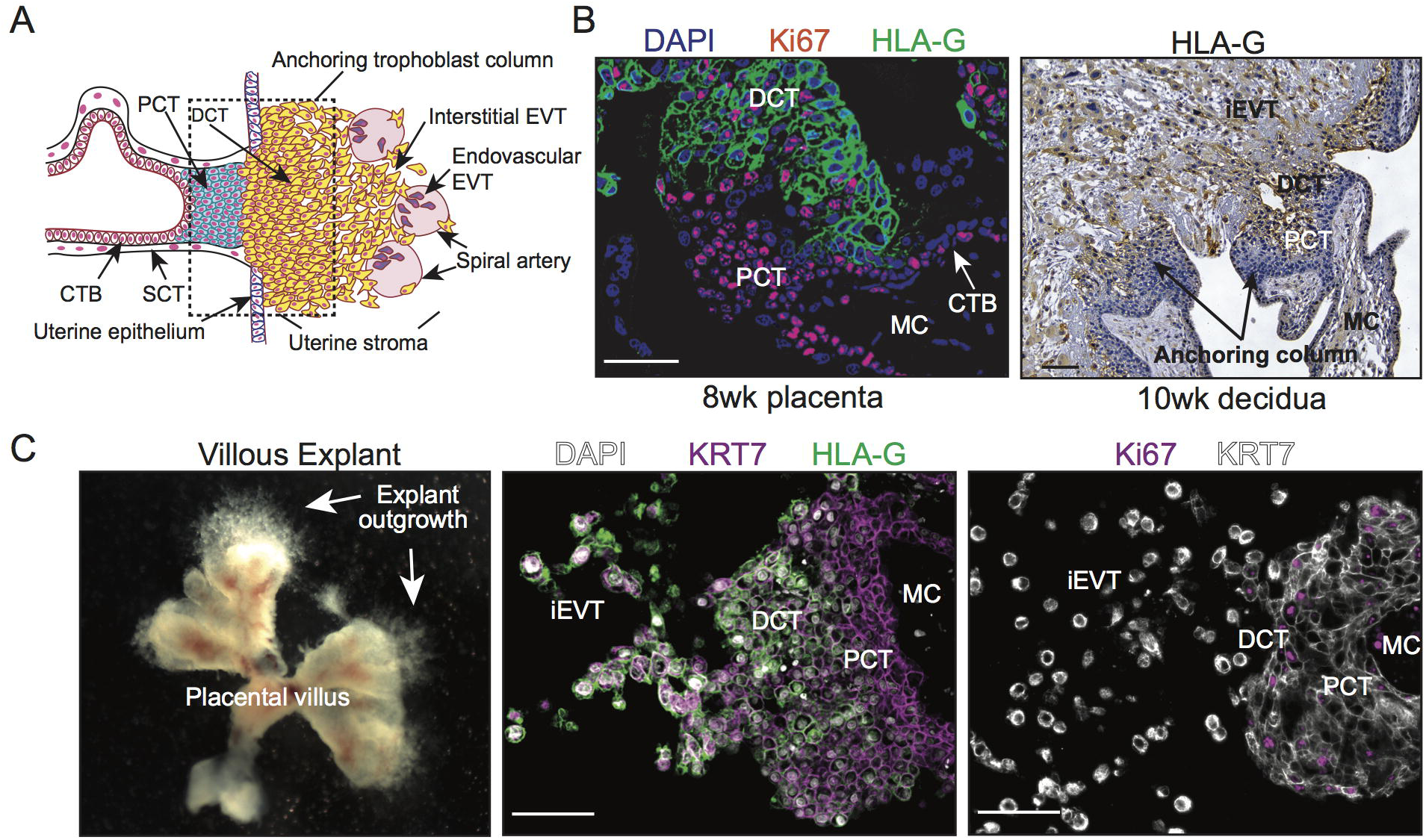
Establishment of a human placental villous explant model. **(A)** Schematic illustration showing trophoblast subtypes within placental villus, anchoring column, and maternal uterine tissue. Depicted are villous cytotrophoblasts (CTB), syncytiotrophoblast (SCT), proximal and distal column trophoblast (PCT & DCT), and invasive interstitial and endovascular EVT. **(B)** Immunofluorescence and immunohistochemistry images showing Ki67 (red) and HLA-G (green, brown) expression in human first trimester placental (8 weeks’ gestation) and decidual (10 weeks’ gestation) tissues. Shown are villous cytotrophoblasts (CTB), proximal and distal column trophoblasts (PCT & DCT), interstitial EVT (iEVT), and the mesenchymal core (MC). Bar = 100 μm. **(C)** Images showing gross villous explant establishment and outgrowth as well as localization of KRT7 (magenta; white), HLA-G (green), and Ki67 (magenta) to specific subtypes of trophoblasts within explant columns. Nuclei are shown via DAPI staining (white). Bar = 100 μm.

Chorionic villi harvested from early first trimester placentae (n=8; 5-8 weeks’ gestation) were imbedded into Matrigel matrix and allowed to establish for 24 hr at 5% oxygen (Figure 2A); this level of oxygen represents a relative “normoxic” condition for early placental development (Jauniaux et al., 2001). Characteristics (age, gestational age, BMI, smoking status) of each patient who donated their placenta for explant culture are listed in Supplemental Table 1. Following this, explants derived from the same placenta were transferred into one of three conditions for an additional 48 hr representing either hypoxic (1%, ∼10 mmHg O_2_), normoxic (5%, ∼35 mmHg O_2_), or hyperoxic (20%, ∼141 mmHg O_2_) environments relative to the first trimester of pregnancy (Figure 2A). Within all oxygen conditions, explant outgrowth was observed (Figure 2B, 2C). However, column outgrowth was most pronounced in 1% and 5% oxygen, where outgrowth area in both of these low oxygen conditions was significantly greater than outgrowth observed in 20% oxygen (Figure 2C, 2D). While column outgrowth in 5% oxygen was overall less variable and trended on producing smaller columns than explants cultured in 1% oxygen, there was no statistical difference between outgrowth in 1% and 5% oxygen (Figure 2D). In summary, trophoblast column outgrowth was potentiated by low oxygen, whereas exposure to atmospheric oxygen (20% oxygen) blunted column outgrowth.

**Figure 2.**
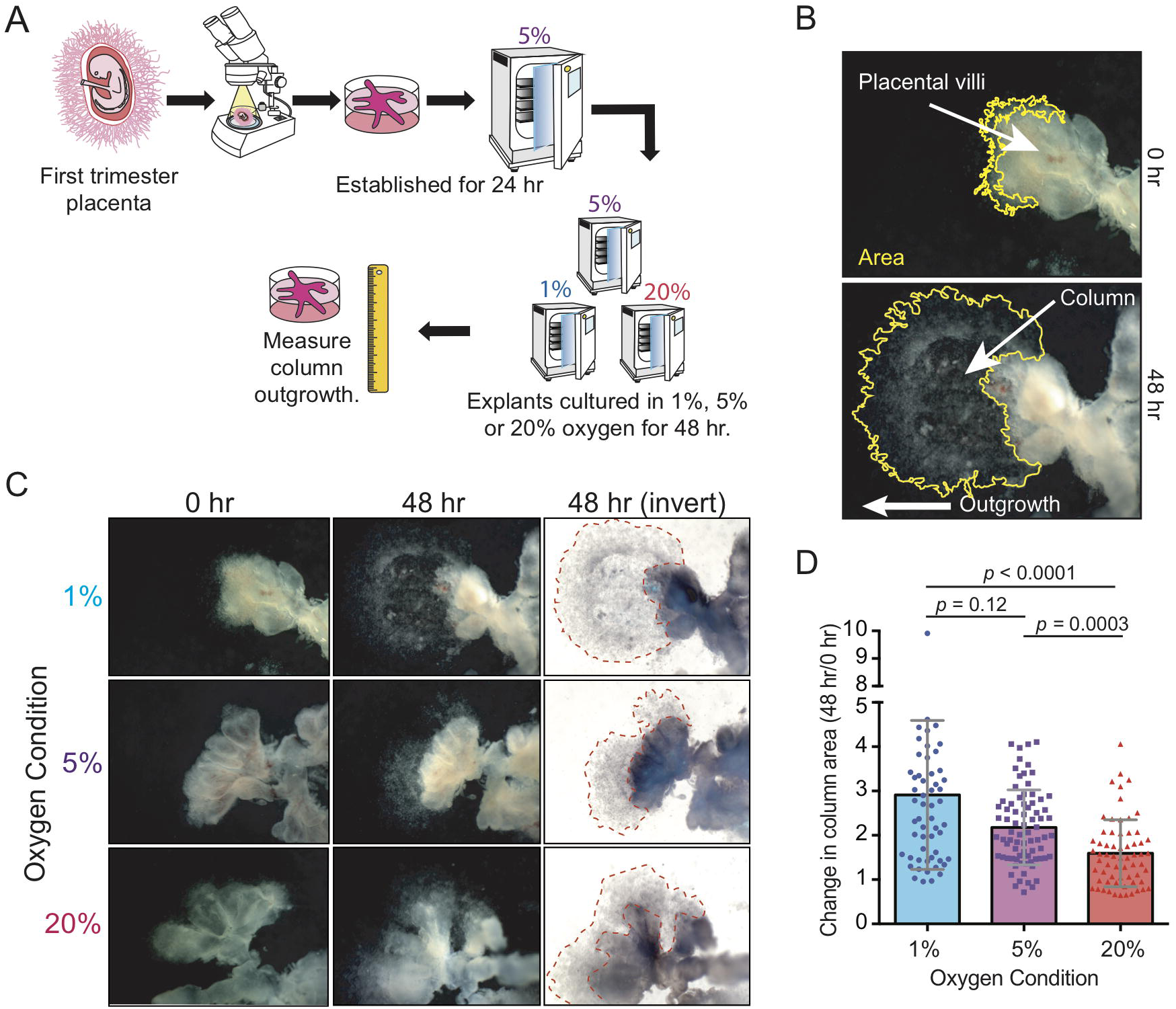
Exposure to low oxygen promotes trophoblast column outgrowth. **(A)** Schematic illustration depicting the experimental approach to establishing and culturing placental explants in 1%, 5%, and 20% oxygen. **(B)** Representative images showing how explant column outgrowth area is measured at 0 hr and 48 hr of culture. The direction of explant outgrowth/invasion is also indicated. **(C)** Representative images of villous explants (n=8 distinct placentae; multiple explants per placenta) cultured in 1%, 5%, and 20% oxygen. Shown within the inverted (invert) image is the area of column outgrowth at 48 hr of culture. **(D)** Bar with scatter plots showing the fold-change of column outgrowth between 48 hr and 0 hr of exposure to 1%, 5%, or 20% oxygen. Median values and standard deviations are shown. Statistical analyses between groups were performed using ANOVA and two-tailed Dunn’s post-test; significance considered *p* < 0.05.

### Explant exposure to low or high levels of oxygen generate distinct transcriptomic signatures

To gain mechanistic insight into how hypoxic, physiologically normal, and high levels of oxygen modulate trophoblast column outgrowth, global gene expression was analysed in placental explant cultures using gene microarrays. For this experiment, placental explants from 5 unique placentae (n=5; 5-7 weeks’ gestation) were established as previously described except that cultures were maintained in their respective oxygen condition (1%, 5%, or 20%) for 24 hr prior to RNA isolation in order to capture molecular signatures central to column formation. Importantly, RNA was extracted from only column trophoblasts and Matrigel-invading EVT; chorionic villi of explants were carefully micro-dissected away from columns and EVT following an approach described within Bilban *et al* (Bilban et al., 2009) (Figure 3A). Following standard probe filtering, and normalization (Supplemental Figure 1), differential gene expression (DGE) analysis of explant trophoblasts was performed. Using a false discover rate (FDR) < 0.05, we identified many differences in gene expression between 1% versus 20% (293 genes up-regulated; 685 genes down-regulated) and 5% versus 20% oxygen conditions (363 genes up-regulated; 406 genes down-regulated) (Figure 3B; Supplemental Table 2). DGE analysis between 1% and 5% oxygen did not identify differentially expressed genes (Figure 3B). Principal component analysis (PCA) shows clustering of explants cultured in hypoxic, normoxic, and hyperoxic conditions (Figure 3C). PCA sample clustering showed explant trophoblasts cultured in 1% and 5% oxygen generally clustered closer together, save for two 1% oxygen explant outliers; one clustered amongst 20% oxygen samples while the other clustered separately to all other samples (Figure 3C). Despite these two explants being identified as significant outliers using the Silhouette coefficient (Barghash and Arslan, 2016), we opted to retain them for the remainder of our analyses as we could not confidently ascribe outlier classification due to technical or batch-related artifacts.

**Figure 3.**
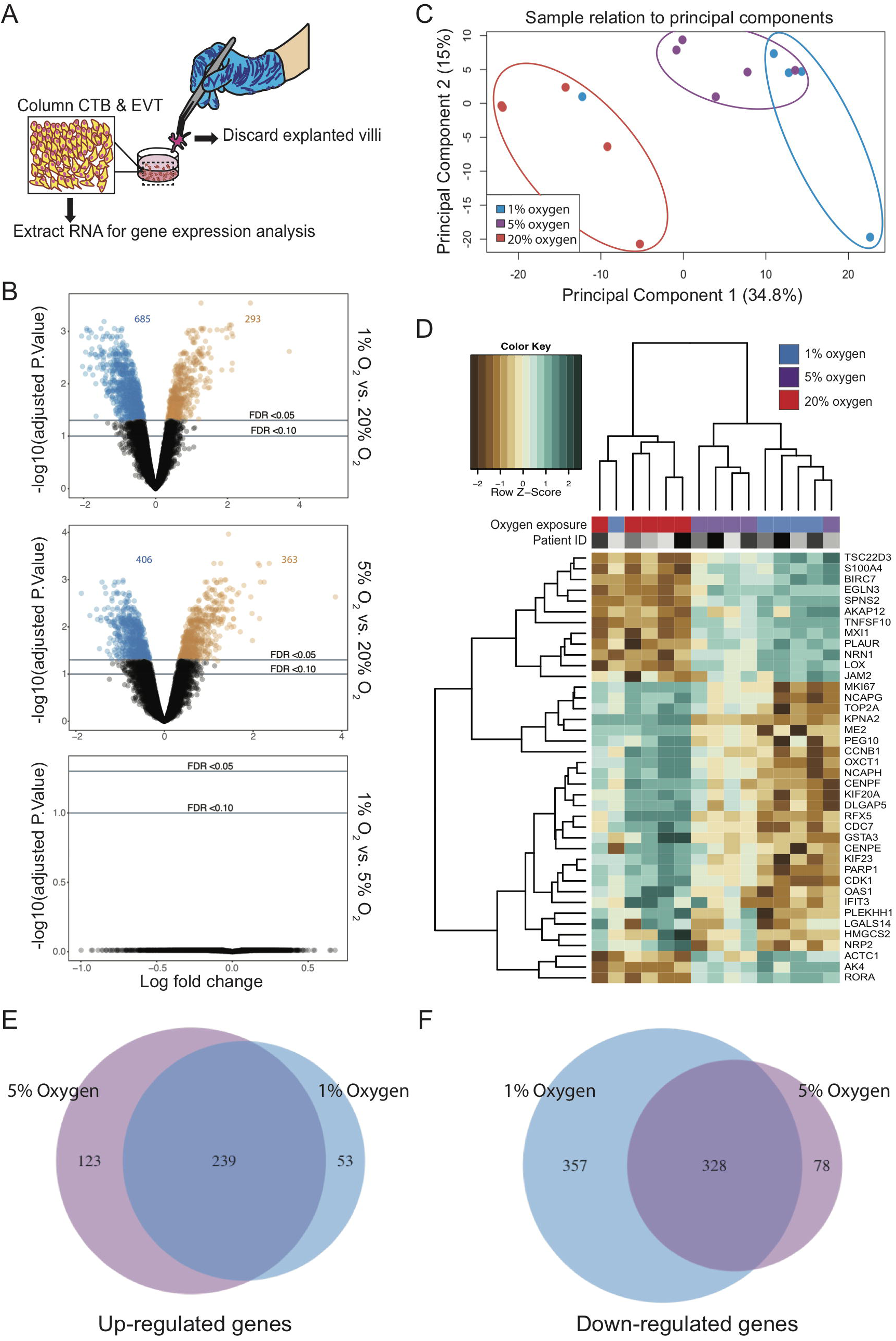
Comparison of global gene expression patterns between trophoblast columns exposed to 1%, 5%, and 20% oxygen. **(A)** Schematic illustration highlighting the removal of placental villi and retention of column trophoblasts and invasive EVT for gene expression analysis. **(B)** Volcano plots showing individual gene-targeting probes differentially expressed between 1% and 20%, 5% and 20%, and 1% and 5% oxygen cultures. X-axis: coefficients from linear model in log2 scale; y axis: negative log base 10 of the false discovery rate (FDR). Black circles indicate probes with an FDR > 0.05; blue and orange circles indicate under-expressed and over-expressed probes with an FDR < 0.05. **(C)** PCA of 1% (blue), 5% (purple), and 20% (red) oxygen cultured explants. **(D)** Dendrogram depicts hierarchical clustering of 1%, 5%, and 20% oxygen cultured column trophoblasts using Z-scores from the top 40 (DEGS from 1% vs 20% comparisons) genes; n=5 per oxygen group. Within the heatmap, brown = low expression; white = mid-level expression; green = high expression. For each sample, oxygen condition (1% = blue; 5% = purple; 20% = red) is indicated, as is the unique placental sample/patient ID used to generate the explant (indicated by shade of grey). Venn diagram showing the number of shared and unique genes (FDR < 0.05) **(E)** up-regulated and **(F)** down-regulated in 1% and 5% oxygen-cultured explants compared to 20% oxygen cultures.

Hierarchical clustering of the 15 column trophoblast samples segregated samples into two statistically significant clusters (*sigclust*, p<0.05): a 20% oxygen dominated cluster and a cluster comprised of column trophoblasts cultured in 1% and 5% oxygen (Figure 3D). A gene heat-map of the top 40 differentially expressed genes (top 20 differentially expressed genes in 1%; top 20 differentially expressed genes in 20%; FDR< 0.05, ranked by FC) highlights gene patterns across 1%, 5%, and 20% oxygen cultures (Figure 3D). In both 1% and 5% oxygen conditions, the top hits identified by global DGE analysis included genes associated with hypoxia (*EGLN3, RORA*), cell-matrix interaction/re-structuring (*LOX, JAM2, EGLN3, PLAUR*), and gene transcription regulation (*MXI1, TSC22D3, RORA*). In contrast, the most highly expressed genes in explants exposed to 20% oxygen were exclusive to pro-mitotic/proliferative processes (*MKI67, KIF20A, KIF23, PEG10, CDK1, NCAPG, NCAPH, TOP2A, CDC7*), indicating that column trophoblasts cultured in 20% oxygen possess a proliferative phenotype.

While stringent DGE parameters did not identify differentially expressed genes between 1% and 5% oxygen culture conditions, we did identify 123 and 53 unique genes to be up-regulated in 5% and 1% oxygen cultures compared to samples cultured in 20% oxygen (Figure 3E). Further, 78 and 357 genes were shown to be uniquely down-regulated in 5% and 1% oxygen cultured explants compared to 20% cultures (Figure 3F). Gene ontology (GO) pathway analysis of the above 1% and 5% signatures indicated that column trophoblasts exposed to 1% oxygen show enrichment for processes favoring hypoxia-related- and oxidative stress-signaling (Supplemental Figure 2). Conversely, column trophoblasts cultured in 5% oxygen showed enrichment of biological pathways linked to nucleotide biosynthesis and metabolism, indicating that column trophoblasts in 1% and 5% oxygen do exhibit underlying molecular differences potentially contributing to trophoblast column development (Supplemental Figure 2).

### Differing oxygen levels drive distinct molecular programs in trophoblast columns

To broadly examine how transcriptomic differences within column trophoblasts exposed to hypoxic, normoxic, and hyperoxic conditions relate to differences in molecular pathways, gene signatures determined by DGE analysis (FDR < 0.05; fold-change > 1.5; Supplemental Table 2) were used to identify pathways enriched in explant column trophoblasts cultured in 1%, 5%, and 20% oxygen. Unsurprisingly, pathways in explants cultured in 1% oxygen (293 genes; clusterProfiler) showed enrichment for multiple pathways and molecular processes specific to hypoxia (Figure 4A). Additionally, the 1% oxygen signature also showed enrichment for pathways related to extracellular matrix (ECM) structure and organization, steroid hormone responses, and hydroxyproline metabolism (Figure 4A). Similar to the 1% oxygen pathway readouts, the 5% oxygen signature (363 genes) showed enrichment for genes specific to ECM composition, response to hypoxia, and response to steroid hormones, but also showed enrichment of pathways related to bone development and viral entry into cells (Supplemental Figure 3). By contrast, 20% oxygen (685 genes) showed enrichment of pathways and cellular processes related to organelle fission, nuclear division, chromosome segregation, mitotic nuclear division, and DNA packaging and replication, all of which link to heightened cell cycle activity and proliferation (Figure 4B).

**Figure 4.**
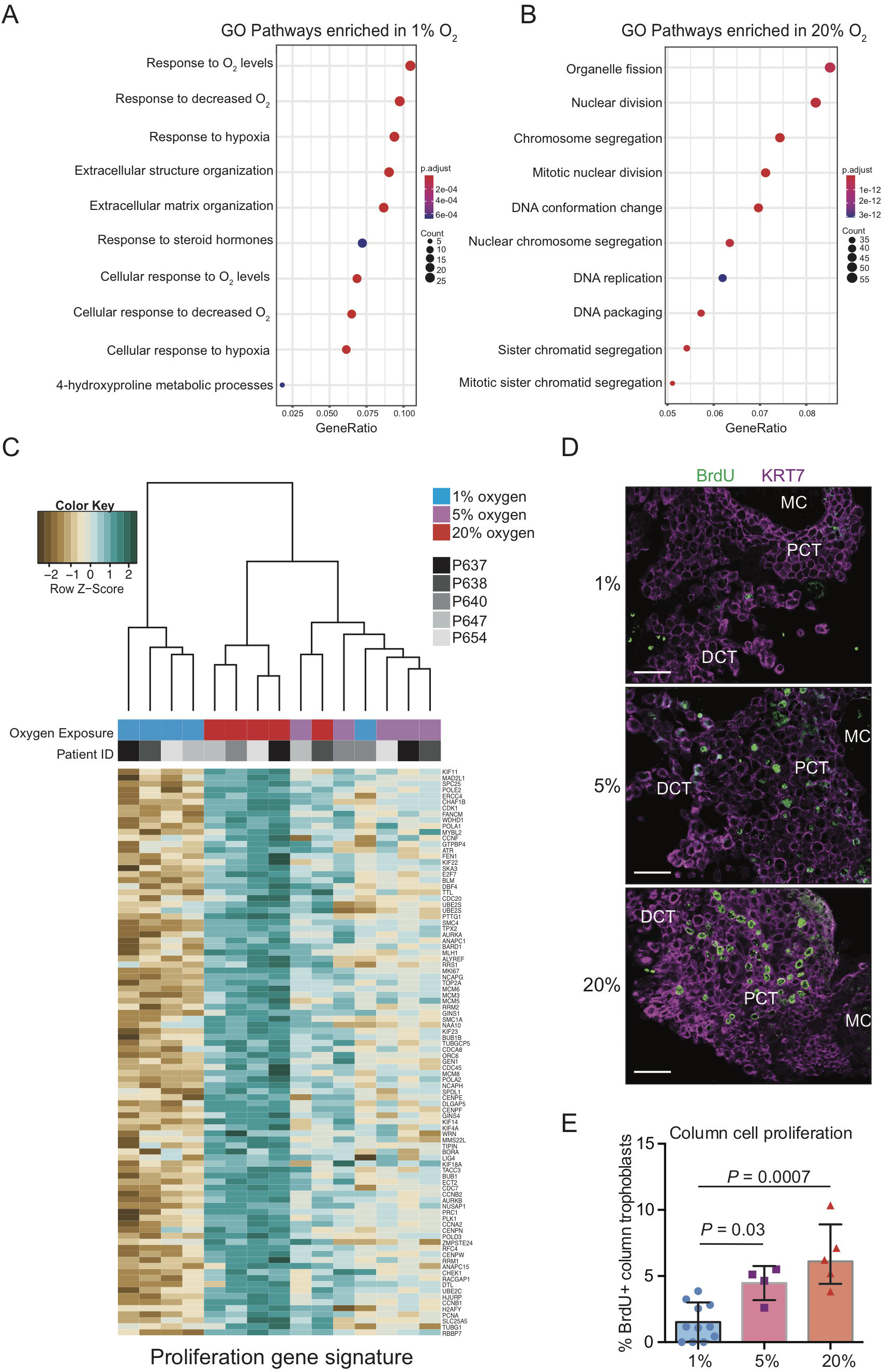
Placental explant exposure to atmospheric oxygen promotes column trophoblast proliferation. Gene ontology (GO) pathway analyses of DGE genes indicating the top 10 up-regulated pathways in explants cultures in **(A)** 1% oxygen and **(B)** 20% oxygen. Adjusted *p* values (Bonferroni) and the number of genes identified in each pathway category is represented beside each plot. **(C)** Dendrogram depicts hierarchical clustering of 1%, 5%, and 20% oxygen cultured column trophoblasts using a proliferation-specific GO signature (Nuclear Division; 93 genes). Within the heatmap, brown = low expression; white = mid-level expression; green = high expression. For each sample, oxygen condition (1% = blue 5% = purple; 20% = red) is indicated, as is the unique placental sample/patient ID used to generate the explant (indicated by shade of grey). **(D)** Representative immunofluorescence images of BrdU signal (green) within placental explant columns cultured in 1%, 5%, or 20% oxygen. Trophoblasts are identified by KRT7 signal (magenta). Bar = 100 μm. **(E)** Bar/scatter plots show quantification of BrdU incorporation into column trophoblasts cultured in 1%, 5%, or 20% oxygen. Median values are shown and statistical analyses between groups were performed using ANOVA and two-tailed Tukey post-test; significance *p* < 0.05.

Using the Mitotic Nuclear Division, Chromosome Segregation, and DNA Replication curated GO gene signatures enriched within 20% column trophoblasts (93 genes; Supplemental Table 3), explant samples were subjected to hierarchical clustering and visualized by gene heat-map (Figure 4C). Notably, samples segregated into two groups: One group defined by 1% oxygen samples (4/5 1% oxygen samples), and the other group consisting of mostly of 5% and 20% oxygen samples (10/10 5%/20% samples) (Figure 4C). This later branch was further divided into two sub-branches, one enriched by 5% oxygen samples also containing a 1% oxygen sample outlier, and the other sub-branch made up entirely of 20% oxygen column trophoblasts (Figure 4C). Interestingly, gene heat-map expression intensities suggest a step-wise increase in expression of pro-mitotic/proliferative genes in column trophoblasts exposed to increasing levels of oxygen (Figure 4C). To verify if an increase in exposure to oxygen tension translates into increased proliferation, a BrdU pulse-chase was performed on a separate cohort of placental explants (n=3) cultured in 1%, 5%, and 20% oxygen. In support of the gene array data, little/no evidence of cell proliferation within explant columns was observed in 1% oxygen cultures following a 4 hr chase (Figure 4D, 4E). However, explants cultured in 5% oxygen showed a significant increase in BrdU incorporation within column trophoblasts compared to 1% oxygen columns, and an even greater level of BrdU positivity was measured within 20% oxygen columns (20% versus 1%) (Figure 4D, 4E). Though a trend for greater proliferation in explant columns cultured in 20% oxygen compared to 5% oxygen was observed, this difference was not significant (Figure 4E). Overall, our findings suggest that explant column trophoblasts cultured in low oxygen up-regulate molecular processes related to hypoxia/HIF1A signalling and ECM organization/remodeling, while column trophoblasts exposed to hyper-physiological 20% oxygen adopt a predominantly pro-proliferative phenotype.

### Low oxygen promotes EVT differentiation

Differentiation of trophoblasts along the EVT pathway is in part defined by the exiting of PCT located at the base of anchoring columns from the cell cycle (Velicky et al., 2018). Moreover, as column trophoblasts located at distal portions of columns acquire pro-invasive EVT-like characteristics, molecular pathways related to ECM-remodeling and protease functions are accordingly up-regulated (Davies et al., 2016). Our observation that low oxygen (1% & 5%) promotes transcriptional signatures linked to cell-ECM interaction and protease-ECM remodeling, while 20% oxygen promotes proliferation, suggests that exposure to low oxygen drives, while high oxygen restrains EVT differentiation.

To gain insight into how differing levels of oxygen affect column trophoblast differentiation, explant column trophoblasts were subjected to hierarchical cluster analysis using a signature of differentially expressed genes derived from a list of 47 trophoblast-related genes curated from recent high-dimensional data and differentiation studies focused on trophoblast biology (Supplemental Figure 4; Supplemental Table 4) (Bilban et al., 2009; Davies et al., 2016; Haider et al., 2018; Lee et al., 2018; Turco et al., 2018). This list includes genes associated with trophoblast lineage (*TFAP2A, KLF5, GATA3*), trophoblast pluripotency (*CDX2*), villous cytotrophoblast (CTB) state (*EGFR, SPINT1, ITGA6, PEG10, TEAD4, TP63*), EVT state (*HLA-G, HTRA1, LAIR2, FLT-1, ERBB2, ADAM12, AMAM19, MYC, ITGA5, TEAD2*), syncytiotrophoblast (SCT) state (*GCM1, CGA, ERVW1, ERVFRD-1, ENDOU*), and genes commonly used to identify proliferative CTB and PCT (*MK167, CCNA2, NOTCH1*). From this curated list, 14 genes were differentially expressed between 1%, 5%, and 20% oxygen cultured explants (Supplemental Figure 4A). Notably, this small signature was sufficient to segregate samples into two main groups: One group consisted almost entirely of column trophoblasts cultured in 20% oxygen (save for one 1% oxygen outlier), while the other group was made up of a mix of 1% and 5% oxygen cultured samples (Supplemental Figure 4A). Genes enriched within low-oxygen cultured explants included trophoblast lineage-related transcription factors (*GATA3*, *KLF5*, *TFAP2A*), genes related to ECM remodeling (*TIMP1*, *ADAM12, ADAM19*), genes related to CTB specification (*CDH1*, *EGFR*), and genes linked with the EVT sub-lineage (*ADAM12*, *FLT1*, *ITGA5, MYC*) (Supplemental Figure 4A). Samples that grouped primarily as 20% oxygen cultured explants showed enrichment for genes specific to proliferative CTB and PCT (*CCNA2*, *MKI67*). Notably, the imprinted paternally expressed gene, *PEG10*, was also highly expressed within 20% oxygen column trophoblasts (Supplemental Figure 4A).

Examination of PEG10 localization within first trimester placental villi (n=3 placentae; 6-10 weeks’ gestation) by immunofluorescence microscopy (IF) showed that PEG10 preferentially localizes to CTB (Supplemental Figure 4B). IF localization of PEG10 within a new cohort of placental explants (established from n=5 placentae) revealed that PEG10 signal, similar to the signal observed in placental villi, is broadly localized to CTB in explants exposed to all three oxygen culture conditions (Supplemental Figure 4C). However, within 20% oxygen explants, bright PEG10 signal was also observed within multi-layered PCT; this column-specific signal was minimal/absent in 1% and 5% oxygen cultures (Supplemental Figure 4C).

To more closely examine how global gene expression changes identified within low and high oxygen exposed explants relate to trophoblast differentiation, we examined the expression of the top fifteen up-regulated genes in column trophoblasts cultured in 1% and 20% oxygen (FDR < 0.05; fold-change > 2) within a recently reported first trimester placenta single cell transcriptomic dataset (Vento-Tormo et al., 2018). Using this dataset we focused exclusively on the 5 subtypes of trophoblasts that were described (14,366 trophoblasts from 5 individual placentae): CTB, proliferative CTB, SCT, proliferative EVT (likely PCT), and EVT (likely an admixture of DCT and invasive EVT) (Figure 5A) (Vento-Tormo et al., 2018). Within UMAP-directed cell clusters, the specificity of each trophoblast sub-lineage/type is shown by expression levels of *EGFR* (CTB), *ERVFRD-1* (syncytin-2; SCT), *HLA-G* (column trophoblast & EVT), and *MKI67* (proliferating trophoblast) (Figure 5B). A heat-map projection shows the pattern of gene expression of trophoblast lineage and subtype-specific genes (Pan trophoblast: *KRT7, TFAP2A, GATA3, KLF5*; CTB: *EGFR, SPINT1, TP63*; proliferative CTB and PCT: *MKI67, CCNA2*; SCT: *ENDOU, ERVFRD-1*; EVT: *HLA-G, ITGA5, ERBB2*), and the fifteen top up-regulated genes in 20% (*OAS1, IFIT3, DLGAP5, GSTA3, CDK1, OXCT1, CENPF, KIF20A, PARP1, TOP2A, PLEKHH1, RFX5, NCPAH, KPNA2, NRP2*) and 1% oxygen conditions (*TNFSF10, LOX, AKAP12, ACTC1, BIRC7, SPNS2, S100A4, JAM2, MXI1, EGLN3, RORA, PLAUR, AK4, TSC22D3, NRN1*) (Figure 5C). Interestingly, genes up-regulated within 1% oxygen-cultured columns showed alignment with proliferative EVT, and this relationship was even greater with EVT (Figure 5C). In contrast, the top genes identified within 20% oxygen columns aligned predominately with proliferative CTB and proliferative EVT (Figure 5C).

**Figure 5.**
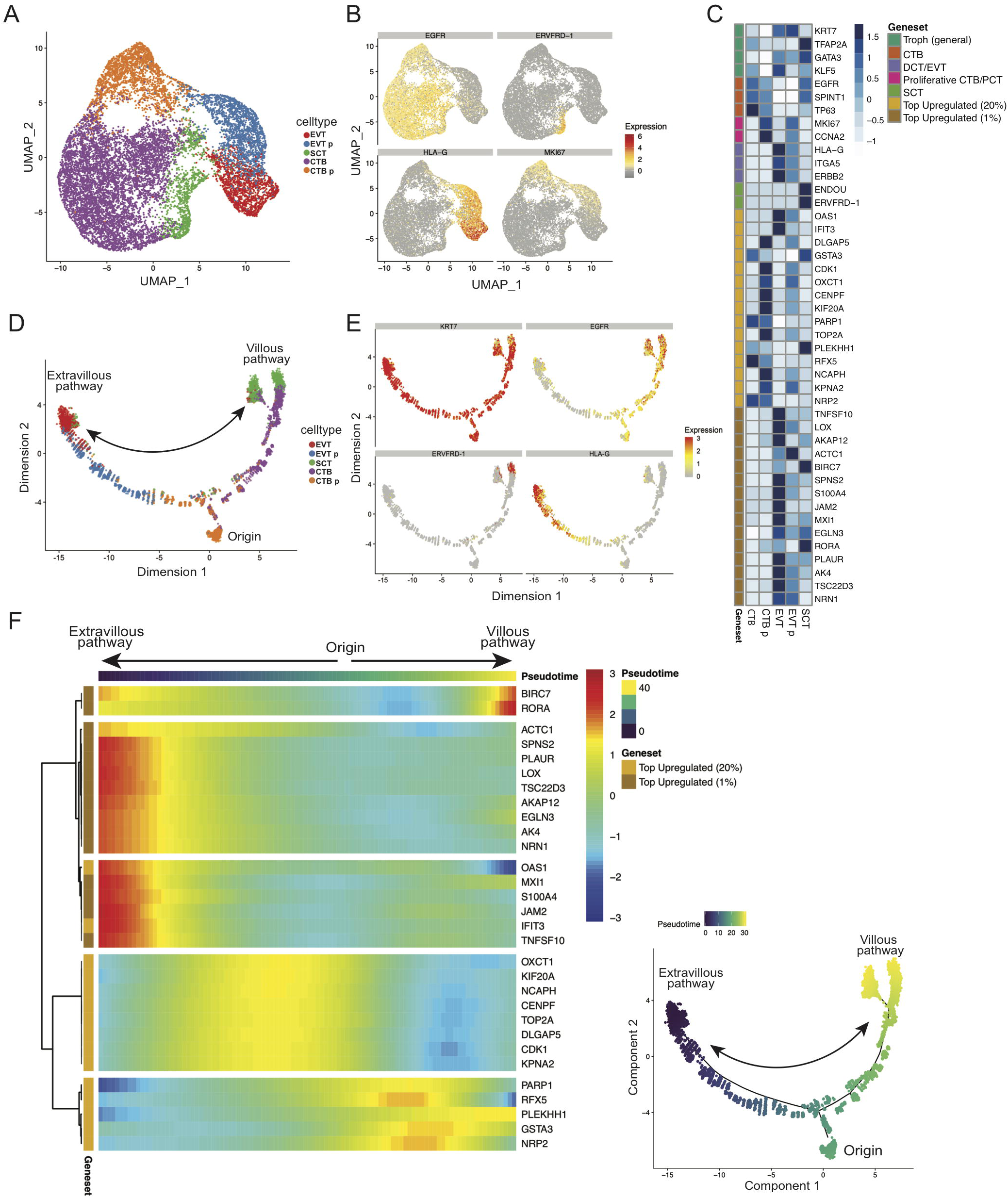
Low oxygen drives column cell differentiation along the EVT pathway. **(A)** Previously published scRNAseq data (Vento-Tormo et al., 2018) for 14,366 trophoblast cells from first trimester placentae (n=5) were selected for investigating the cell-specificity of our list of identified oxygen-associated gene expression changes. Uniform Manifold Approximation and Projection (UMAP) (Becht et al., 2018) was used to visualize and cluster the cells after subsetting from other cell types in the published dataset. Cells are labelled according to their previous characterization (Vento-Tormo et al., 2018). **(B)** UMAP clusters cells by gene expression of canonical trophoblast marker genes. **(C)** Heatmap of top 15 genes upregulated in 1% and 20% oxygen. Also shown are genes aligning with general trophoblasts, villous cytotrophoblasts (CTB), proximal column trophoblast (PCT), and distal column trophoblast (DCT)/extravillous trophoblast (EVT). Geneset expression patterns are compared to averaged gene expression levels within single-cell informed cell types (CTB, proliferative CTB, EVT, proliferative EVT, SCT). **(D)** Pseudotime analysis was applied using Monocle 2 (Qiu et al., 2017; Trapnell et al., 2014) to visualize gene expression across trophoblast differentiation. Two lineage trajectories were identified corresponding to the extravillous pathway and villous pathway. A cell state of origin is also shown. **(E)** The inferred trajectory resulted into two distinct endpoints: one branch leading to cells highly expressing SCT markers (e.g. ERVFRD-1), and another leading to cells highly expressing EVT markers (HLA-G). **(F)** A heatmap was constructed using inferred pseudotime and the top 15 upregulated genes in 1% and in 20% oxygen conditions. Pseudotime was ordered such that the left and right ends represent the EVT and SCT endpoints. Hierarchical clustering was applied to the genes (ordered along the rows) and separated into 5 clusters.

Pseudotime trajectory analysis using a gene signature derived from the top 1000 variable genes reproduced a lineage trajectory similar to that reported in Vento-Tormo *et al* (Vento-Tormo et al., 2018), where a predicted cell origin state was identified as well as the two differentiation trajectories aligning with the villous and extravillous pathways (Figure 5D). Correlating the expression of genes specific to distinct states of trophoblast differentiation (i.e. *EGFR, HLA-G, ERVFRD-1*) with the pseudotime trajectory confirmed the accuracy of the pseudotime modeling, where *EGFR* aligned to cells committed to the villous pathway, *HLA-G* to a maturing EVT, and *ERVFRD-1* to a terminally differentiated SCT (Figure 5E). Correlating trophoblast lineage trajectory with the top up-regulated 1% and 20% oxygen genes showed that genes enriched within 20% oxygen aligned to both the villous and extravillous pathway trajectories (Figure 5F). Notably, high-oxygen genes aligned closely with cell states linked to proliferative CTB (*PARP1, RFX5, PLEKHH1, GSTA3, NRP2*) and proliferative PCT (*OXCT1, KIF20A, NCAPH, CENPF, TOP2A, DLGAP5, CDK1, KPNA2*) (Figure 5F). By contrast, low oxygen enriched genes primarily aligned with the tail-end of the extravillous pathway (i.e. *TNFSF10, LOX, SPNS2, S100A4, JAM2, PLAUR*) (Figure 5F). Together, these data suggest that column trophoblasts exposed to low oxygen adopt transcriptomic signatures that are reflective of EVT, while column trophoblasts cultured in 20% oxygen express genes that align predominately with proliferating CTB and PCT.

### *LOX* expression and activity is potentiated by low oxygen

Our finding that low oxygen drives column outgrowth and potentiates the expression of genes linked with the EVT phenotype suggests that genes highly expressed within low oxygen columns may in part contribute to EVT differentiation and trophoblast column formation. Rank ordering of up-regulated genes by fold-change in both 1% and 5% oxygen column trophoblasts identified multiple conserved genes between the two oxygen conditions (Supplemental Table 2). Notably, *LOX*, the gene encoding lysyl oxidase, a copper-dependent enzyme that catalyses collagen and elastin crosslinking, was the number 2-ranked gene in both 1% and 5% oxygen cultured explants. Specifically, *LOX* expression was 6.2- and 5.1-fold higher in 1% and 5% oxygen cultures compared to 20% oxygen explants (Figure 6A; Supplemental Table 2). While previous work has identified a role for elevated *LOX* expression in promoting tumor cell metastasis (Cox et al., 2015; Di Stefano et al., 2016), the role of LOX in placental trophoblast column biology and trophoblast differentiation along the EVT pathway has not been described.

**Figure 6.**
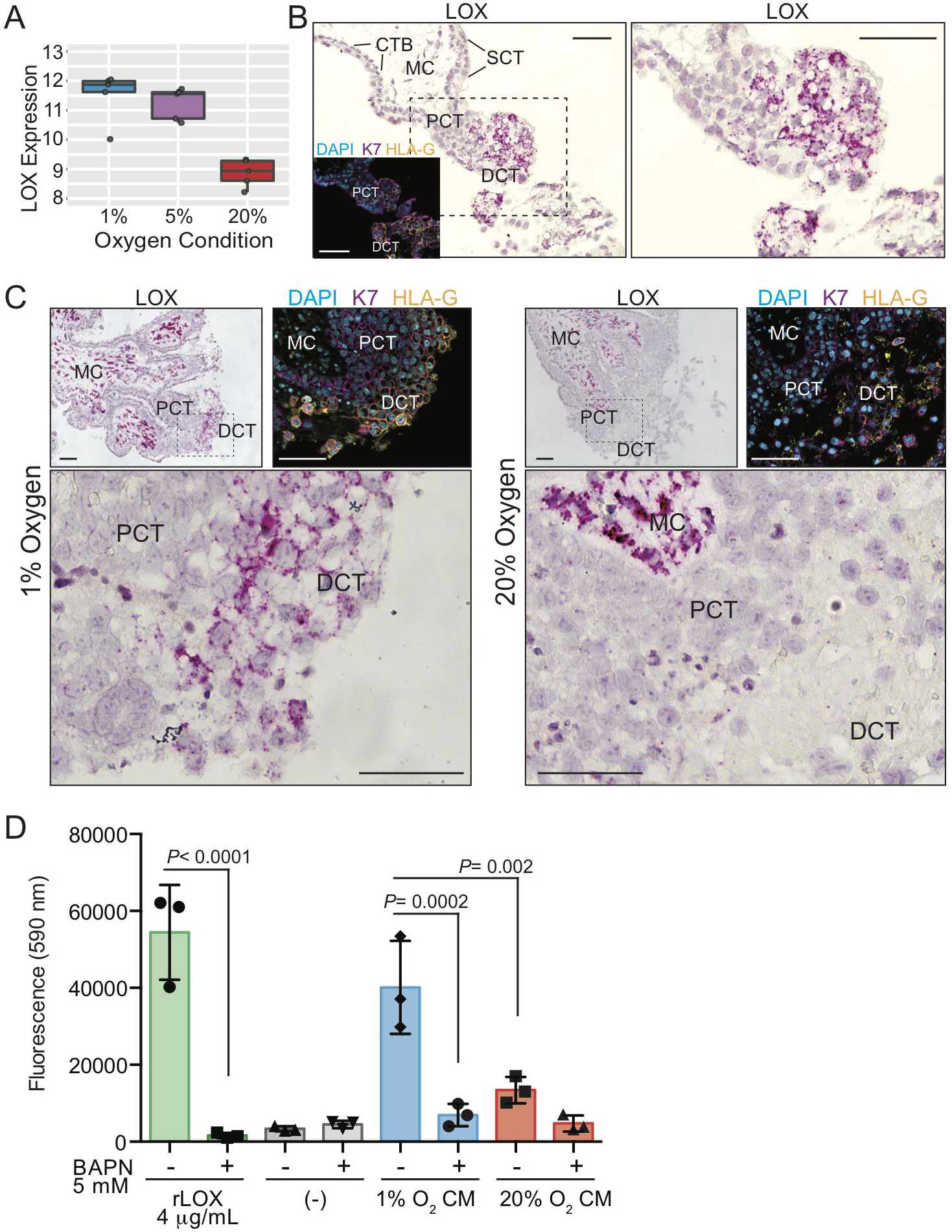
Low oxygen exposure promotes column-specific expression of *LOX*. **(A)** Gene array box-plot expression levels of *LOX* mRNA in placental explants cultured in 1%, 5%, and 20% oxygen. **(B)** Representative image showing *LOX* mRNA transcript *in situ* localization (dark pink signal) within a first trimester placental villus. The hashed box outlines the enlarged region to the right. Shown are annotations of trophoblast/placental subtypes; Mesenchymal core (MC), cytotrophoblast (CTB), syncytiotrophoblast (SCT), proximal column trophoblast (PCT), distal column trophoblast (DCT). Shown as an inset is an immunofluorescent image depicting nuclei (DAPI; blue), keratin-7 (K7; magenta), and HLA-G (orange) localization within a serial section of the same placental villus. Bars = 100 μm. **(C)** Representative images of *LOX* mRNA localization within placental explants cultured in 1% or 20% oxygen. Specific trophoblast subtypes are indicated as above, and the hashed box corresponds to the enlarged area shown below. Also shown are insets of immunofluorescent images depicting nuclei (DAPI; blue), keratin-7 (K7; magenta), and HLA-G (orange) localization within corresponding serial sectioned regions of the explant. Bars = 100 μm. **(D)** Bar/scatter plots show LOX activity levels within conditioned media (CM) of placental explants cultured in either 1% or 20% oxygen in the presence/absence of the LOX inhibitor BAPN (5 mM). Recombinant active lysyl oxidase (rLOX) served as a positive control whereas explant culture media alone served as a negative control. Activity corresponds to level of fluorescence intensity (590 nm). Median values are shown and statistical analyses between groups were performed using ANOVA and two-tailed Tukey post-test; significance *p* <0.05.

As an initial step to examine the importance of LOX in anchoring column biology, *LOX* mRNA *in situ* hybridization within first trimester placental villi (n=3; 6-8 weeks’ gestation) was performed. RNAscope *in situ* hybridization showed specific and intense *LOX* localization to cells within the mesenchymal core of placental villi and to trophoblasts within anchoring columns (Figure 6B). Little/no *LOX* signal was detected in CTB or SCT (Figure 6B). RNAscope analysis of *LOX* within placental explants cultured in 1% and 20% oxygen supported the gene array finding that *LOX* expression was elevated in column trophoblasts exposed to low oxygen (Figure 6C). While we were unable to verify elevated LOX protein expression in low oxygen-cultured placental explants via IF microscopy due to non-specific antibody signal, LOX enzymatic activity, measured in conditioned media (CM) generated by placental explants cultured in 1% or 20% oxygen, showed that activity was significantly higher in 1% cultures (Figure 6D). Use of the LOX inhibitor, β-aminopropionitrile (BAPN), demonstrated LOX-substrate specificity, while recombinant active LOX served as a positive control (Figure 6D). Taken together, these findings show that LOX expression and activity is elevated in column trophoblasts cultured in low oxygen. Further, *LOX’s* preferential *in vivo* expression within the trophoblast anchoring column combined with its involvement in promoting tumor cell metastasis, suggests that LOX may also play a role in controlling trophoblast column outgrowth and/or EVT differentiation.

### Impairment of LOX restrains column outgrowth

To test the function of LOX in column outgrowth, Matrigel-imbedded placental explants cultured in 1% oxygen (n=3) were cultured in either control explant media or media containing the competitive LOX inhibitor BAPN. The effectiveness of BAPN in inhibiting LOX activity was measured as before by examining the ability of endogenous LOX in explant CM to oxidize substrate (Figure 7A). Conditioned media harvested from control explants showed LOX activity levels slightly below levels measured in recombinant LOX positive control reactions, but significantly higher than explant media alone (Figure 7A). Importantly, treatment of explants with BAPN significantly blunted LOX activity, though activity was not completely blocked (Figure 7A). Importantly, placental explant treatment with BAPN led to a significant two-fold impairment in column outgrowth (Figure 7B, 7C). Taken together, these results suggest that LOX expression in developing trophoblast columns promotes column outgrowth and associates with an EVT phenotype.

**Figure 7.**
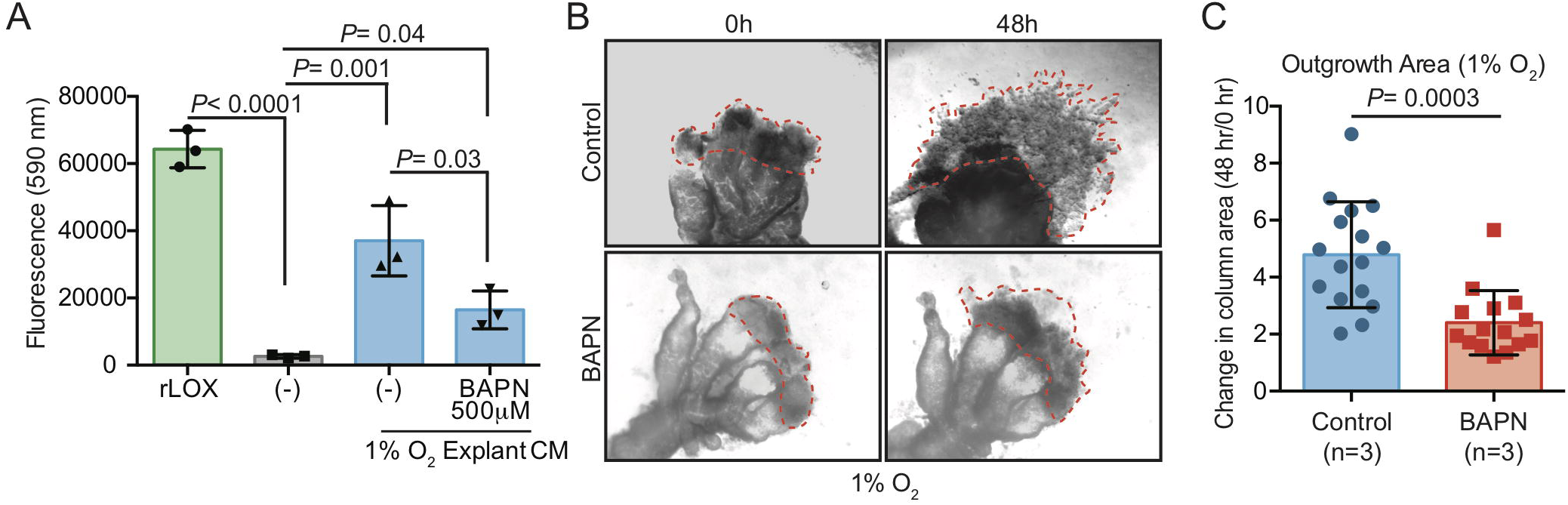
Inhibition of lysyl oxidase (LOX) dampens column outgrowth. **(A)** Bar/scatter plots show LOX activity levels within conditioned media (CM) of placental explants cultured in 1% oxygen in the presence/absence of BAPN (500 μM). Recombinant active lysyl oxidase (rLOX) served as a positive with explant culture media alone served as a negative control. Activity corresponds to level of fluorescence intensity (590 nm). Median values are shown and statistical analyses between groups were performed using ANOVA and two-tailed Tukey post-test. **(B)** Representative images of villous explants (n=3 distinct placentae; multiple explants per placenta) cultured in 1% oxygen and in the presence/absence of BAPN (500 μM). Column outgrowth is shown at 0hr and 48 hr of culture. **(C)** Box plots showing the fold-change of column outgrowth between 48 hr and 0 hr following treatment with BAPN. Median values are shown and statistical analyses between groups were performed using ANOVA and two-tailed Tukey post-test; significance *p* <0.05.

## DISCUSSION

Here we describe how exposure to different levels of oxygen differentially affect trophoblast column outgrowth and global gene expression. We provide evidence that exposure to low oxygen results in overall increases in column outgrowth accompanied by gene expression signatures that align with an EVT phenotype. By contrast, gene signatures in high oxygen-cultured column trophoblasts define a role for elevated oxygen in maintaining column growth through cell proliferation. In both hypoxic and normoxic conditions, we identify the gene *LOX*, as one of the most highly up-regulated genes within explant columns. We show that *LOX* expression associates with EVT lineage trajectory and demonstrate that impairment of LOX activity blunts column outgrowth. Together, this work supports a role for low oxygen in potentiating the EVT pathway. Moreover, this work also identifies novel oxygen-sensitive molecular processes that likely play roles in anchoring column formation during human placental development.

The role of oxygen in controlling column formation and the EVT pathway is controversial. Previous studies have shown that exposure of placental explants to low oxygen (i.e. 2% to 3% oxygen) promotes column expansion and outgrowth, where outgrowth is primarily attributed to HIF1A-directed cell proliferation (Caniggia et al., 2000; Genbacev et al., 1997). Consistent with this, evidence exists that CTB exposure to low oxygen restrains trophoblast progression along the EVT pathway (Caniggia et al., 2000; Genbacev et al., 1997; Lash et al., 2006). By contrast, rodents and rodent-derived trophoblast stem cells engineered to lack hypoxia-sensing machinery (i.e. *ARNT*, *HIF1A*, and/or *EPSA1* null) fail to differentiate into trophoblast lineages of the labyrinth zone and into trophoblast giant cells, trophoblast populations akin to invasive EVT in humans (Chakraborty et al., 2016; Chakraborty et al., 2011; Cowden Dahl et al., 2005; Gultice et al., 2009; Maltepe et al., 2005). In support of these observations, low oxygen was shown to potentiate an invasive phenotype in human primary trophoblasts with an accompaniment in the expression of EVT-associated genes HLA-G and α5 integrin, and the up-regulation of pro-migratory integrin-linked kinase signaling (Horii et al., 2016; Robins et al., 2007; Wakeland et al., 2017). Our findings overall align with these later studies that suggest low oxygen promotes differentiation along the EVT pathway. Notably, we show that low oxygen drives expression of EVT-related genes and signatures that align with EVT lineage trajectory. We show that low oxygen (1% and 5% oxygen) induces expression of hallmark EVT genes like *ITGA5*, *ADAM12*, and *FLT1*, as well as the transcription factors *KLF5* and *GATA3* that are preferentially expressed by EVT. Importantly, the use of cell lineage trajectory modeling using single cell RNA sequencing data also provides evidence that low oxygen promotes a cell state consistent with the EVT lineage. These later observations are in line with the association of low levels of oxygen within the intervillous space and anchoring column formation and interstitial EVT infiltration into decidual mucosa in early pregnancy. However, the relationship between relative placental hypoxia and inadequate placentation in aberrant pregnancy conditions like preeclampsia (Farrell et al., 2019) suggests that the impact of oxygen on trophoblast biology may differ according to stage of development.

Our finding that high levels of oxygen promote column trophoblast proliferation is inconsistent with previous studies investigating the role of hypoxia in anchoring column biology (Caniggia et al., 2000; Genbacev et al., 1997). However, in agreement with our findings, James *et al* reported that explant columns cultured in 8% oxygen have increased trophoblast cellularity compared with columns exposed to 1.5% oxygen (James et al., 2006). Moreover, there seems to be consistent agreement that exposure to low oxygen leads to greater column outgrowth, where oxygen tension levels between 1-5% consistently generate larger columns than atmospheric oxygen conditions (Caniggia et al., 2000; James et al., 2006). However, dissecting how enhanced column outgrowth is achieved, i.e. via greater cell proliferation within the column or through increased trophoblast migration and invasion, is still not completely resolved. Indeed, while our data suggests that low oxygen promotes an EVT phenotype and enhances molecular pathways related to cell-ECM remodeling, we did not directly examine if low oxygen affects EVT invasion. The stark differences on the effect of oxygen levels in regulating trophoblast proliferation and EVT differentiation within explant columns between various studies is difficult to explain, but may stem from differences in media composition, the matrix substratum used in explant cultures, subtle variations in oxygen tension, and the gestational age of placental tissues/cells used for establishing explant cultures. Future studies will need to specifically re-examine how differing levels of oxygen impact column cell proliferation.

*PEG10*, a paternally expressed and maternally imprinted gene, was identified as a top up-regulated gene within 20% oxygen column trophoblasts. Previous work has shown a critical role for Peg10 in mouse placental development, where mice deficient in Peg10 show severe fetal growth restriction and fetal demise by E10.5, in addition to defects in labyrinth and spongiotrophoblast development (Ono et al., 2006). Intriguingly, a partial loss in *Peg10* imprinting leading to *Peg10* over-expression also associates with impaired labyrinth formation (Koppes et al., 2015). These findings suggest that a fine balance in Peg10 expression is required for proper placental development. Notably, in humans, dysregulation of PEG10 associates with certain pregnancy disorders, including pre-eclampsia, gestational hypertension, molar pregnancy, and spontaneous miscarriage (Dória et al., 2010; Liang et al., 2014; Rahat et al., 2017). Our finding that *PEG10* expression and cell proliferation in trophoblast columns associates with exposure to high oxygen is consistent with studies showing a role for PEG10 in promoting tumor cell survival and growth (Bang et al., 2015; Ishii et al., 2017; Peng et al., 2017). Recent findings also provide evidence that PEG10 promotes trophoblast cell line proliferation (Abed et al., 2019). Given that PEG10 specifically localizes to CTB of floating villi and to subsets of trophoblasts within proximal regions of anchoring columns suggests that PEG10 may play roles in column establishment by promoting progenitor expansion. Further work is needed to dissect the role of PEG10 in column formation and progression along the EVT pathway.

Our finding that *LOX* expression was consistently a gene highly up-regulated in both 1% and 5% column trophoblasts compared to those cultured in 20% oxygen indicates that LOX-related processes are important in placental development and trophoblast column biology. While a role of LOX in column formation has not been previously reported, the importance of LOX in tumorigenesis is known, where elevated LOX associates with numerous types of cancers and LOX activity promotes tumor cell metastasis (Cox et al., 2015; Di Stefano et al., 2016). Further, a recent report does provide evidence that LOX expression promotes cell invasion of a trophoblast cell line (Xu et al., 2019). This later finding is consistent with the association of LOX expression and the EVT phenotype. Aldehydes produced by LOX-directed oxidation of lysine residues within collagen and elastin facilitate collagen/elastin cross-linking and stability, which in turn provide a structural lattice for cell movement (Kim et al., 2014). That *LOX* expression is greatest within both proximal and distal column trophoblasts is interesting, as PCT are not considered to be migratory. Nonetheless, it is likely that creating an appropriate substratum scaffold via LOX-directed collagen crosslinking may contribute to column stability and provide a platform for EVT outgrowth. How or if LOX affects anchoring column establishment or EVT differentiation is unknown, but future studies utilizing newly-derived trophoblast organoids (Haider et al., 2018; Turco et al., 2018) may allow for deep mechanistic examination of LOX and the EVT pathway.

In summary, the extravillous pathway is controlled by multiple intrinsic as well as extrinsic factors, including level of oxygen. We provide evidence that supports a role for low oxygen levels in promoting the differentiation of trophoblasts along the EVT pathway. This finding establishes insight into critical developmental events during placentation that occur in early pregnancy. Further, these findings may also provide a foundation for understanding cellular and molecular processes contributing to conditions linked to aberrant placentation.

## MATERIALS AND METHODS

### Patient recruitment and tissue collection

Decidual and placental tissues were obtained with approval from the Research Ethics Board on the use of human subjects, University of British Columbia (H13-00640). All samples were collected from women (19 to 39 years of age) providing written informed consent undergoing elective terminations of pregnancy at British Columbia’s Women’s Hospital, Vancouver, Canada. First trimester decidual (*N*=1) and placental tissues (*N*=30) were collected from participating women (gestational ages ranging from 5–12 weeks) having confirmed viable pregnancies by ultrasound-measured fetal heartbeat. The decidual tissue sample was selected based on the presence of a smooth uterine epithelial layer and a textured thick spongy underlayer. Patient clinical characteristics i.e. height and weight were additionally obtained to calculate body mass index (BMI: kg/m^2^) and all consenting women provided self-reported information via questionnaire to having first hand exposure to cigarette smoke, and are summarized in Supplemental Table 1.

### Placental villous explant assay

*Ex vivo* placental villous cultures were established as described in (Aghababaei et al., 2014; De Luca et al., 2017; Perdu et al., 2016). Briefly, placental villi from 5-8 week old gestation placentas (n=8) obtained from patients undergoing elective termination of pregnancy were dissected, washed in cold PBS, and imbedded into Millicell cell culture inserts (0.4 μm pores, 12mm diameter. EMD Millipore, Billerica, MA) containing 200 μl of growth-factor-reduced Phenol-red free Matrigel (BD Biosciences, San Diego, CA). Explants, containing 400 μl DMEM/F12 1:1 (200 mM L-glutamine) in the outer chamber, were allowed to establish overnight in a humidified 37 °C trigas incubator at 5% oxygen, 5% CO_2_. Following 24 hr of culture, explants were cultured in 200 μl DMEM/F12 1:1 media and placed into 1%, 5%, or 20% oxygen incubators for either an additional 24 hr (gene expression analyses) or 48 hr (explant outgrowth measurements). All explant media were supplemented with penicillin/streptomycin and antimycotic solution (Thermofisher Scientific, Waltham, MA) Growing explants were imaged at indicated times using a Nikon SMZ 7454T triocular dissecting microscope (Minato, Japan) outfitted with a digital camera. EVT outgrowths were measured by ImageJ software. Fold-change in outgrowth was determined by dividing the mean column area at 48 hr into the mean area at 0 hr.

### Immunofluorescence, RNAScope, and immunohistochemistry microscopy

*Immunofluorescence:* Placental villi (6-12 weeks gestation; n=5) or placental explants (derived from n=11 placentae) were fixed in 2% paraformaldehyde overnight at 4 °C. Tissues were paraffin embedded and sectioned at 6 μm onto glass slides. Immunofluorescence was performed as described elsewhere (Aghababaei et al., 2015). Briefly, cells or placental tissues underwent antigen retrieval by heating slides in a microwave for 5 × 2 minute intervals in a citrate buffer (pH 6.0). Sections were incubated with sodium borohydride for 5 minutes, RT, followed by Triton X-100 permeabilization for 5 minutes, RT. Slides were blocked in 5% normal goat serum/0.1% saponin for 1h, RT, and incubated with combinations of the indicated antibodies overnight, 4 °C: Anti-HLA-G (1:100, 4H84, Exbio, Vestec, Czech Republic); anti-cytokeratin 7, mouse monoclonal IgG (1:50, RCK105, Santa Cruz Biotechnology, Dallas, TX); anti-cytokeratin 7, rabbit monoclonal IgG (1:50, SP52, Ventana Medical Systems, Oro Valley, AZ); anti-Ki67 (1:75, SP6, Thermo Fisher Scientific); anti-LOX (1:100, NB100-2527, NovusBio, Centennial, CO); anti-PEG10 (1:100, 4C10A7, NovusBio); anti-BrdU (1:1000, Bu20a, Cell Signalling Technology, Danvers, MA). Following overnight incubation, sections and coverslips were washed with PBS and incubated with Alexa Fluor goat anti rabbit-488/-568 and goat anti mouse-488/-568 conjugated secondary antibodies (Life Technologies, Carlsbad, CA) for 1 hr at RT and washed in PBS, and mounted using ProLong Gold mounting media containing DAPI (Life Technologies).

Slides were imaged with an AxioObserver inverted microscope (Car Zeiss, Jena, Germany) using 20X Plan-Apochromat/0.80NA or 40X Plan-Apochromat oil/1.4NA objectives (Carl Zeiss). An ApoTome .2 structured illumination device (Carl Zeiss) set at full Z-stack mode and 5 phase images was used for image acquisition. For quantification of BrdU signal in column trophoblasts, 2-4 cell columns per explant (n=9; obtained using a 20X objective) were used to calculate values; BrdU+ cell proportions were calculated by BrdU/KRT7+ cells per column; BrdU fluorescence intensity thresholds were used to calculate BrdU proportions within Matrigel explant columns. Images were obtained using an Axiocam 506 monochrome digital camera and processed and analyzed using ZenPro software (Carl Zeiss).

*Immunohistochemistry:* Decidua (10 weeks’ gestation; n=1) from a first trimester pregnancy was fixed in 2% paraformaldehyde for 24 hr at 4°C, paraffin imbedded, and serially sectioned at 6µm onto glass slides. Heat-induced antigen retrieval was performed using sodium citrate (10mM, pH 6.0) followed by quenching endogenous peroxidases with 3% hydrogen peroxide for 30 minutes at RT. Sections were then permeabilized with 0.2% Triton-X-100 for 5 minutes at RT. Serum block was performed with 5% BSA in tris-buffered saline with 0.05% Tween 20 (TBST). Sections were then incubated with mouse monoclonal HLA-G (1:100, clone 4H84, ExBio) diluted in TBST overnight at 4°C. Following overnight incubation, sections were incubated with Envision+ Dual Link Mouse/Rabbit HRP-linked secondary antibody (DAKO, Santa Clara, CA) for 1 hr at RT. IgG1 isotype controls and secondary antibody-only negative controls were performed to confirm antibody specificity. Staining was developed via 3,3’-diaminobenzidine (DAB) chromogen (DAB Substrate Kit, Thermo Scientific), counterstained in Modified Harris Hematoxylin Solution (Sigma-Aldrich, St. Louis, MO) and coverslips mounted with Entellan mounting medium (Electron Microscopy Sciences, Hatfield, PA).

*RNAScope:* RNA *in situ* hybridization was performed using RNAscope^®^ 2.5 HD Assay-RED (Advanced Cell Diagnostics, Newark, CA) following the manufacturer’s instructions (Wang *et al*., 2012). Briefly, placenta (n = 3; 6-8 weeks’ gestation) and Matrigel-imbedded placental explants (n=15; 5 explants per oxygen condition) from first trimester pregnancies were fixed overnight at 4°C in 4% paraformaldehyde and paraffin-embedded. Tissue sections were serially sectioned at 6 µm, and following deparaffinization, antigen retrieval was performed using RNAscope^®^ Target Retrieval Reagent (95°C for 15 minutes) and ACD Protease Plus Reagent (40°C for 30 minutes). RNAScope probes targeting LOX (Hs-LOX-C2; Ref. 415941-C2) and negative control (Probe-dapB; Ref. 310043) were incubated on sections for 2hr at 40°C, and following this, the RNAscope signal was amplified over 6 rounds of ACD AMP 1-6 incubation and application of RED-A and RED-B at a ratio of 1 volume of RED-B to 60 volumes of RED-A for 10 minutes at RT. Selected samples were counterstained with 50% Hematoxylin I for 2 minutes at RT and all samples were mounted using EcoMount (Biocare Medical, Pacheco, CA).

### RNA purification

Total RNA was prepared from column and invasive EVT cells using TRIzol reagent (Life Technologies, Carlsbad, CA) followed by RNeasy MinElute Cleanup (Qiagen, Hilden, Germany) and DNase treatment (Life Technologies) according to the manufacturer’s instructions. Care was taken so that explants were exposed to atmospheric oxygen no longer than 2 minutes during microscopic dissection and separation of placental villi away from columns prior to the addition of TRIzol. RNA purity was confirmed using a NanoDrop Spectrophotometer (Thermo Fisher Scientific) and by running RNA samples on an Agilent 2100 Bioanalyzer (Agilent Technologies, Santa Clara, CA). Only RNA samples having an RNA Integrity Number (RIN) > 8.0 were used.

### Microarray hybridization, gene array data preprocessing, gene expression analysis

Total RNA samples extracted from explant columns were sent to Génome Québec Innovation Centre (McGill University, Montréal, Canada) for RNA quantification. Briefly, RNA samples were prepared for transcriptome profiling using the GeneChip ™ Pico Reagent Kit (Thermo Fisher Scientific) as per manufacturer’s protocol. Samples were run on the Clariom™ S Human Array to measure gene expression at >20,000 genes in the human genome (Affymetrix, Santa Clara, CA). Raw data generated from the arrays were read into R statistical software (version 3.5.1) with the Bioconductor *oligo* package to convert raw Affymetrix CEL files into an expression matrix of intensity values. The expression data was background corrected, Quantile normalized, and log-transformed. A total of 13,787 control, duplicated, non-annotated or low intensity probes were filtered out of the data, leaving 13,402 probes for further analysis. Pre-processing was monitored at each step by Principal Component Analysis (PCA) and linear modelling. Principal component analysis was performed by the `svd()` function from the `sva` package in R. Linear modelling was conducted using the R package *limma*. Probe-wise variances were shrunk using empirical Bayes with the `eBayes` function, followed by FDR adjustment for multiple testing (Benjamini and Hochberg, 1995). Differentially expressed genes were defined based on an FDR < 0.05. Enrichment of pathways were identified and annotated using the *clusterProfiler* package in Bioconductor (Carvalho and Irizarry, 2010). Volcano plots were generate using the *ggplot2* package for RStudio. Cluster analysis of sample relations based on principal components was generated using the `plotSampleRelation` function for the *Lumi* package. A hierarchical cluster analysis was conducted using Euclidean distances on the top 40 differentially expressed genes, selected by FDR <0.05 and ranked by fold-change > 1.5.

### Single cell RNA-seq data analysis

Processed droplet-based and Smart-seq2 single cell RNA sequencing data was obtained from public repositories (ArrayExpress experiment codes: E-MTAB-6701 and E-MTAB-6678) (Vento-Tormo et al., 2018). Cells belonging to clusters from trophoblast lineages (n=14,366) were merged from droplet-based and Smart-seq2 data, and corrected for batch-effects using canonical correlation analysis implemented in the R package *Seurat* (Butler et al., 2018). Pseudotime trajectory modelling was conducted using the *monocle 2* R package (Qiu et al., 2017; Trapnell et al., 2014) under the recommended unsupervised procedure called “`dpFeature`”. Briefly, the first 10 principal components on log-normalized expression data were used to construct a TSNE projection, upon which density-peak clustering determined 13 number of clusters, using parameters rho = 2, delta = 10. The top 1000 differentially expressed genes between these clusters were then used for ordering the cells. Visualization of oxygen concentration-dependent genes were visualized along the inferred pesudotime-ordered branches using the R function `monocle::plot_genes_branched_heatmap` with the following settings: the number of clusters k = 5 for clustering the genes, and default parameters for all else.

### LOX activity assay and LOX inhibition

Measurement of endogenous LOX activity in placental explant conditioned media was performed following a modified protocol described in Wiel et al (Wiel et al., 2013). Briefly, 600 μl of conditioned media from placental explants cultured in triplicate from either 1%, 5%, or 20% oxygen conditions was pooled, concentrated 12-fold using 7.5 kDa exclusion Amicon Millipore concentration columns (Millipore, Burlington, MA), and snap-frozen in liquid nitrogen. LOX activity in 15 μl of concentrated conditioned media was determined using the Amplex Red H_2_O_2_ detection kit following the manufacture’s instructions (Life Technologies, Carlsbad, CA). This assay is based on the ability of endogenous LOX to oxidize 10 mM DAP (a LOX substrate) in the presence of 0.5 U/ml horseradish peroxidase; the reaction is incubated at 37 °C for 30 minutes. Oxidation of Amplex Red generates a fluorescence signal measurable at 560nm/590nm excitation/emission wavelengths and was detected using a fluorescence plate reader (BMG Labtech, Ortenberg, Germany) using a 96-well format. As a positive control, 10 μg/ml recombinant active LOX (MyBioscource.com, San Diego, CA) was separately incubated with DAP substrate. LOX activity/specificity was determined by co-incubating reactions with 5 mM of BAPN. For endogenous inhibition of LOX within explant cultures, following 24 hr of explant establishment at 37 °C, 5% CO2, 5% O2 in a humidified tri-gas incubator, explant media was replaced with media containing 500 μM of BAPN and explant cultures were placed into 1% or 20% O_2_ culture conditions for an additional 48 hr of culture prior to being imaged/measured.

### BrdU incorporation assay

Pulse-chase labeling with BrdU (Sigma-Aldrich) was conducted on a verification cohort of Matrigel embedded placental explants (n=3 distinct placentae, 4-11 columns per oxygen condition). Explants were established in 5% oxygen for 24 hr followed by 24 hr of culture in either 1%, 5% or 20% oxygen. After 48 hr of culture, explants were exposed to a 4-hr pulse with culture media containing 10μM of BrdU. Following 4 hr of labelling, explants were washed in PBS and fixed in 4% PFA overnight. Explants were paraffin embedded and sectioned for immunofluorescence microscopy. Immunofluorescent staining with anti-BrdU antibody (Bu20a, Sigma-Aldrich) with the addition of a 30-minute incubation in 2M hydrochloric acid between permeabilization and sodium borohydride steps.

### Statistical analysis

Data are reported as median values with standard deviations. All calculations were carried out using GraphPad Prism software (San Diego, CA). For single comparisons, Mann-Whitney non-parametric unpaired t-tests were performed. For multiple comparisons, one-way Kruskal-Wallis ANOVA followed by Dunn’s multiple comparison test was performed on explant outgrowth data as outgrowth in 1% oxygen was not normally distributed. One-way ANOVA followed by Tukey post-test were performed for all other multiple comparisons. The differences were accepted as significant at *P* < 0.05. For gene microarray and scRNA-seq statistical analyses, please refer to the Gene Array Data Preprocessing and scRNA-seq analysis sections in methods.

## Abbreviations

BAPN: β-aminopropionitrile
BrdU: Bromodeoxyuridine
CM: Conditioned media
CTB: Cytotrophoblast
DAP: 1,5-diaminopentane
DCT: Distal column trophoblast
DGE: Differential gene expression
ECM: Extracellular matrix
EVT: Extravillous trophoblast
FDR: False discovery rate
GO: Gene ontology
HIF1A: Hypoxia inducible factor 1 A
HLA-G: Human leukocyte antigen G
Hr: Hour
IF: Immunofluorescence
LOX: Lysyl oxidase
PCA: Principal component analysis
PCT: Proximal column trophoblast
PEG10: Paternally expressed gene 10
scRNA-seq: single cell RNA sequencing
SCT: Syncytiotrophoblast
UMAP: Uniform manifold approximation and projection
vCTB: Villous cytotrophoblast

## DATA AVAILABILITY/ACCESSION NUMBER

The GEO accession number for the data reported in this paper is: GSE132421

## AUTHOR CONTRIBUTIONS

AGB designed the research. AGB, JT, VY, JB, HL, and BC performed experiments and analysed data. AGB, JT, VY, and WPR wrote the paper. All authors read and approved the manuscript.

## ACKNOWLEDGEMENTS

The authors extend their sincere gratitude to the hard work of staff at British Columbia’s Women’s Hospital’s CARE Program for recruiting participants to our study, and thank Dr. Megan K. Barker, for her critical readings of the manuscript.

## Supplemental Figure Legends

**Supplemental Figure 1.** Normalization and PCA analysis of placental explant gene microarray data. Average expression values for each explant sample **(A)** before and **(B)** after quantile normalization and filtering/removal of probes. **(C and D)** Principal component analysis (PCA) of raw data, normalized data, and filtered data.

**Supplemental Figure 2.** Pathway analyses of unique gene signatures of 1% and 5% oxygen-cultured explants. **(A)** Top 20 gene pathways upregulated exclusively in 1% oxygen when compared to 20% oxygen. **(B)** Top 20 gene pathways exclusively upregulated in 5% oxygen when compared to 20% oxygen. The size of the dot represents the number of significant genes in the pathway and the colour represents adjusted *P*-values as indicated in the legend.

**Supplemental Figure 3.** Top gene pathways identified in explant columns cultured in 5% oxygen. Top 20 pathways upregulated in 5% oxygen when compared to 20% oxygen. The size of the dot represents the number of significant genes in the pathway and the colour represents adjusted *P*-values as indicated in the legend.

**Supplemental Figure 4.** Low oxygen culture promotes trophoblastic and EVT gene signatures in placental explants. **(A)** Dendrogram depicts hierarchical clustering of 1%, 5%, and 20% oxygen cultured column trophoblasts using the respective 14-gene signature. The heatmap displays expression profiles of trophoblast genes identified as being significantly different between 1%, 5%, and 20% cultures. Within the heatmap, brown = low expression; white = mid-level expression; green = high expression. For each sample, oxygen condition (1% = blue 5% = purple; 20% = red) is indicated, as is the unique placental sample/patient ID used to generate the explant (indicated by shade of grey). Representative immunofluorescence images showing expression of PEG10 (magenta) within **(B)** first trimester placental villi and **(C)** Matrigel-imbedded placental explants cultured in either 1%, 5%, or 20% oxygen. Trophoblast are identified by keratin-7 staining (KRT7; green) and nuclei are stained with DAPI (white). Shown are specific cell types: Syncytiotrophoblast (SCT), villous cytotrophoblasts (CTB), proximal column trophoblast (PCT), distal column trophoblast (DCT), and the mesenchymal core (MC). Bar = 100 μm.

## REFERENCES

Abed, M., Verschueren, E., Budayeva, H., Liu, P., Kirkpatrick, D. S., Reja, R., Kummerfeld, S. K., Webster, J. D., Gierke, S., Reichelt, M., et al. (2019). The Gag protein PEG10 binds to RNA and regulates trophoblast stem cell lineage specification. PLoS ONE 14, e0214110.

Aghababaei, M., Hogg, K., Perdu, S., Robinson, W. P. and Beristain, A. G. (2015). ADAM12-directed ectodomain shedding of E-cadherin potentiates trophoblast fusion. Cell Death Differ. 22, 1970–1984.

Aghababaei, M., Perdu, S., Irvine, K. and Beristain, A. G. (2014). A disintegrin and metalloproteinase 12 (ADAM12) localizes to invasive trophoblast, promotes cell invasion and directs column outgrowth in early placental development. Mol. Hum. Reprod. 20, 235–249.

Avagliano, L., Marconi, A. M., Romagnoli, S. and Bulfamante, G. P. (2012). Abnormal spiral arteries modification in stillbirths: the role of maternal prepregnancy body mass index. 25, 2789–2792.

Bang, H., Ha, S. Y., Hwang, S. H. and Park, C.-K. (2015). Expression of PEG10 Is Associated with Poor Survival and Tumor Recurrence in Hepatocellular Carcinoma. Cancer Res Treat 47, 844–852.

Becht, E., McInnes, L., Healy, J., Dutertre, C.-A., Kwok, I. W. H., Ng, L. G., Ginhoux, F. and Newell, E. W. (2018). Dimensionality reduction for visualizing single-cell data using UMAP. Nat. Biotechnol. 37, 38–44.

Benjamini, Y. and Hochberg, Y. (1995). Controlling the False Discovery Rate: A Practical and Powerful Approach to Multiple Testing. Journal of the Royal Statistical Society: Series B (Methodological) 57, 289–300.

Bilban, M., Haslinger, P., Prast, J., Klinglmüller, F., Woelfel, T., Haider, S., Sachs, A., Otterbein, L. E., Desoye, G., Hiden, U., et al. (2009). Identification of novel trophoblast invasion-related genes: heme oxygenase-1 controls motility via peroxisome proliferator-activated receptor gamma. Endocrinology 150, 1000–1013.

Butler, A., Hoffman, P., Smibert, P., Papalexi, E. and Satija, R. (2018). Integrating single-cell transcriptomic data across different conditions, technologies, and species. Nat. Biotechnol. 36, 411–420.

Caniggia, I., Mostachfi, H., Winter, J., Gassmann, M., Lye, S. J., Kuliszewski, M. and Post, M. (2000). Hypoxia-inducible factor-1 mediates the biological effects of oxygen on human trophoblast differentiation through TGFbeta(3). J. Clin. Invest. 105, 577–587.

Carvalho, B. S. and Irizarry, R. A. (2010). A framework for oligonucleotide microarray preprocessing. Bioinformatics 26, 2363–2367.

Chakraborty, D., Cui, W., Rosario, G. X., Scott, R. L., Dhakal, P., Renaud, S. J., Tachibana, M., Rumi, M. A. K., Mason, C. W., Krieg, A. J., et al. (2016). HIF-KDM3A-MMP12 regulatory circuit ensures trophoblast plasticity and placental adaptations to hypoxia. Proc. Natl. Acad. Sci. U.S.A. 113, E7212–E7221.

Chakraborty, D., Rumi, M. A. K., Konno, T. and Soares, M. J. (2011). Natural killer cells direct hemochorial placentation by regulating hypoxia-inducible factor dependent trophoblast lineage decisions. Proc. Natl. Acad. Sci. U.S.A. 108, 16295–16300.

Cowden Dahl, K. D., Fryer, B. H., Mack, F. A., Compernolle, V., Maltepe, E., Adelman, D. M., Carmeliet, P. and Simon, M. C. (2005). Hypoxia-inducible factors 1alpha and 2alpha regulate trophoblast differentiation. Molecular and Cellular Biology 25, 10479–10491.

Davies, J. E., Pollheimer, J., Yong, H. E. J., Kokkinos, M. I., Kalionis, B., Murthi, P., Davies, J. E., Yong, H. E. J., Kokkinos, M. I., Kalionis, B., et al. (2016). Epithelial-mesenchymal transition during extravillous trophoblast differentiation. Cell Adhesion & Migration 10, 310–321.

De Luca, L. C., Le, H. T., Mara, D. L. and Beristain, A. G. (2017). ADAM28 localizes to HLA-G(+) trophoblasts and promotes column cell outgrowth. Placenta 55, 71–80.

Dória, S., Sousa, M., Fernandes, S., Ramalho, C., Brandão, O., Matias, A., Barros, A. and Carvalho, F. (2010). Gene expression pattern of IGF2, PHLDA2, PEG10 and CDKN1C imprinted genes in spontaneous miscarriages or fetal deaths. Epigenetics 5, 444–450.

Farrell, A., Alahari, S., Ermini, L., Tagliaferro, A., Litvack, M., Post, M. and Caniggia, I. (2019). Faulty oxygen sensing disrupts angiomotin function in trophoblast cell migration and predisposes to preeclampsia. JCI Insight 4.

Fock, V., Plessl, K., Draxler, P., Otti, G. R., Fiala, C., Knöfler, M. and Pollheimer, J. (2015a). Neuregulin-1-mediated ErbB2-ErbB3 signalling protects human trophoblasts against apoptosis to preserve differentiation. J. Cell. Sci. 128, 4306–4316.

Fock, V., Plessl, K., Fuchs, R., Dekan, S., Milla, S. K., Haider, S., Fiala, C., Knöfler, M. and Pollheimer, J. (2015b). Trophoblast subtype-specific EGFR/ERBB4 expression correlates with cell cycle progression and hyperplasia in complete hydatidiform moles. Hum. Reprod. 30, 789–799.

Genbacev, O., Zhou, Y., Ludlow, J. W. and Fisher, S. J. (1997). Regulation of human placental development by oxygen tension. Science 277, 1669–1672.

Gultice, A. D., Kulkarni-Datar, K. and Brown, T. L. (2009). Hypoxia-inducible factor 1alpha (HIF1A) mediates distinct steps of rat trophoblast differentiation in gradient oxygen. Biology of Reproduction 80, 184–193.

Haider, S., Meinhardt, G., Saleh, L., Fiala, C., Pollheimer, J. and Knöfler, M. (2016). Notch1 controls development of the extravillous trophoblast lineage in the human placenta. Proc. Natl. Acad. Sci. U.S.A. 113, E7710–E7719.

Haider, S., Meinhardt, G., Saleh, L., Kunihs, V., Gamperl, M., Kaindl, U., Ellinger, A., Burkard, T. R., Fiala, C., Pollheimer, J., et al. (2018). Self-Renewing Trophoblast Organoids Recapitulate the Developmental Program of the Early Human Placenta. Stem Cell Reports 11, 537–551.

Horii, M., Li, Y., Wakeland, A. K., Pizzo, D. P., Nelson, K. K., Sabatini, K., Laurent, L. C., Liu, Y. and Parast, M. M. (2016). Human pluripotent stem cells as a model of trophoblast differentiation in both normal development and disease. Proc. Natl. Acad. Sci. U.S.A. 113, E3882–91.

Ishii, S., Yamashita, K., Harada, H., Ushiku, H., Tanaka, T., Nishizawa, N., Yokoi, K., Washio, M., Ema, A., Mieno, H., et al. (2017). The H19-PEG10/IGF2BP3 axis promotes gastric cancer progression in patients with high lymph node ratios. Oncotarget 8, 74567–74581.

James, J. L., Stone, P. R. and Chamley, L. W. (2006). The effects of oxygen concentration and gestational age on extravillous trophoblast outgrowth in a human first trimester villous explant model. Human Reproduction 21, 2699–2705.

Jauniaux, E., Watson, A. and Burton, G. (2001). Evaluation of respiratory gases and acid-base gradients in human fetal fluids and uteroplacental tissue between 7 and 16 weeks’ gestation. YMOB 184, 998–1003.

Kabir-Salmani, M., Shiokawa, S., Akimoto, Y., Sakai, K. and Iwashita, M. (2004). The role of alpha(5)beta(1)-integrin in the IGF-I-induced migration of extravillous trophoblast cells during the process of implantation. Molecular Human Reproduction 10, 91–97.

Koppes, E., Himes, K. P. and Chaillet, J. R. (2015). Partial Loss of Genomic Imprinting Reveals Important Roles for Kcnq1 and Peg10 Imprinted Domains in Placental Development. PLoS ONE 10, e0135202.

Lash, G. E., Otun, H. A., Innes, B. A., Bulmer, J. N., Searle, R. F. and Robson, S. C. (2006). Low oxygen concentrations inhibit trophoblast cell invasion from early gestation placental explants via alterations in levels of the urokinase plasminogen activator system. Biol. Reprod. 74, 403–409.

Lee, C. Q. E., Turco, M. Y., Gardner, L., Simons, B. D., Hemberger, M. and Moffett, A. (2018). Integrin α2 marks a niche of trophoblast progenitor cells in first trimester human placenta. Development 145, dev162305.

Liang, X. Y., Chen, X., Jin, Y. Z., Chen, X. O. and Chen, Q. Z. (2014). Expression and significance of the imprinted gene PEG10 in placenta of patients with preeclampsia. Genet. Mol. Res. 13, 10607–10614.

Maltepe, E., Krampitz, G. W., Okazaki, K. M., Red-Horse, K., Mak, W., Simon, M. C. and Fisher, S. J. (2005). Hypoxia-inducible factor-dependent histone deacetylase activity determines stem cell fate in the placenta. Development 132, 3393–3403.

Newby, D., Marks, L., Cousins, F., Duffie, E. and Lyall, F. (2005). Villous explant culture: characterization and evaluation of a model to study trophoblast invasion. Hypertens Pregnancy 24, 75–91.

Ono, R., Nakamura, K., Inoue, K., Naruse, M., Usami, T., Wakisaka-Saito, N., Hino, T., Suzuki-Migishima, R., Ogonuki, N., Miki, H., et al. (2006). Deletion of Peg10, an imprinted gene acquired from a retrotransposon, causes early embryonic lethality. Nat. Genet. 38, 101–106.

Peng, Y.-P., Zhu, Y., Yin, L.-D., Zhang, J.-J., Wei, J.-S., Liu, X., Liu, X.-C., Gao, W.-T., Jiang, K.-R. and Miao, Y. (2017). PEG10 overexpression induced by E2F-1 promotes cell proliferation, migration, and invasion in pancreatic cancer. J. Exp. Clin. Cancer Res. 36, 30.

Perdu, S., Castellana, B., Yooona, K., Chan, K., DeLuca, L. and Beristain, A. G. (2016). Maternal obesity drives functional alterations in uterine NK cells. JCI Insight 1.

Pijnenborg, R., Vercruysse, L. and Carter, A. M. (2011). Deep trophoblast invasion and spiral artery remodelling in the placental bed of the chimpanzee. Placenta 32, 400–408.

Pollheimer, J., Vondra, S., Baltayeva, J., Beristain, A. G. and Knöfler, M. (2018). Regulation of Placental Extravillous Trophoblasts by the Maternal Uterine Environment. Front Immunol 9, 297.

Qiu, X., Mao, Q., Tang, Y., Wang, L., Chawla, R., Pliner, H. A. and Trapnell, C. (2017). Reversed graph embedding resolves complex single-cell trajectories. Nat. Methods 14, 979–982.

Rahat, B., Mahajan, A., Bagga, R., Hamid, A. and Kaur, J. (2017). Epigenetic modifications at DMRs of placental genes are subjected to variations in normal gestation, pathological conditions and folate supplementation. Sci Rep 7, 40774.

Robins, J. C., Heizer, A., Hardiman, A., Hubert, M. and Handwerger, S. (2007). Oxygen tension directs the differentiation pathway of human cytotrophoblast cells. Placenta 28, 1141–1146.

Tilburgs, T., Crespo, Â. C., van der Zwan, A., Rybalov, B., Raj, T., Stranger, B., Gardner, L., Moffett, A. and Strominger, J. L. (2015). Human HLA-G+ extravillous trophoblasts: Immune-activating cells that interact with decidual leukocytes. Proc. Natl. Acad. Sci. U.S.A. 112, 7219–7224.

Trapnell, C., Cacchiarelli, D., Grimsby, J., Pokharel, P., Li, S., Morse, M., Lennon, N. J., Livak, K. J., Mikkelsen, T. S. and Rinn, J. L. (2014). The dynamics and regulators of cell fate decisions are revealed by pseudotemporal ordering of single cells. Nat. Biotechnol. 32, 381–386.

Turco, M. Y., Gardner, L., Kay, R. G., Hamilton, R. S., Prater, M., Hollinshead, M. S., McWhinnie, A., Esposito, L., Fernando, R., Skelton, H., et al. (2018). Trophoblast organoids as a model for maternal-fetal interactions during human placentation. Nature 213, 1.

Velicky, P., Knöfler, M. and Pollheimer, J. (2016). Function and control of human invasive trophoblast subtypes: Intrinsic vs. maternal control. Cell Adhesion & Migration 10, 154–162.

Velicky, P., Meinhardt, G., Plessl, K., Vondra, S., Weiss, T., Haslinger, P., Lendl, T., Aumayr, K., Mairhofer, M., Zhu, X., et al. (2018). Genome amplification and cellular senescence are hallmarks of human placenta development. PLoS Genet. 14, e1007698.

Vento-Tormo, R., Efremova, M., Botting, R. A., Turco, M. Y., Vento-Tormo, M., Meyer, K. B., Park, J.-E., Stephenson, E., Polański, K., Goncalves, A., et al. (2018). Single-cell reconstruction of the early maternal-fetal interface in humans. Nature 563, 347–353.

Wakeland, A. K., Soncin, F., Moretto-Zita, M., Chang, C.-W., Horii, M., Pizzo, D., Nelson, K. K., Laurent, L. C. and Parast, M. M. (2017). Hypoxia Directs Human Extravillous Trophoblast Differentiation in a Hypoxia-Inducible Factor–Dependent Manner. The American Journal of Pathology 187, 767–780.

Wallace, A. E., Fraser, R. and Cartwright, J. E. (2012). Extravillous trophoblast and decidual natural killer cells: a remodelling partnership. Hum. Reprod. Update 18, 458–471.

Wang, F., Flanagan, J., Su, N., Wang, L.-C., Bui, S., Nielson, A., Wu, X., Vo, H.-T., Ma, X.-J. and Luo, Y. (2012). RNAscope: a novel in situ RNA analysis platform for formalin-fixed, paraffin-embedded tissues. J Mol Diagn 14, 22–29.

Wiel, C., Augert, A., Vincent, D. F., Gitenay, D., Vindrieux, D., Le Calvé, B., Arfi, V., Lallet-Daher, H., Reynaud, C., Treilleux, I., et al. (2013). Lysyl oxidase activity regulates oncogenic stress response and tumorigenesis. Cell Death Dis 4, e855–e855.

Xu, X.-H., Jia, Y., Zhou, X., Xie, D., Huang, X., Jia, L., Zhou, Q., Zheng, Q., Zhou, X., Wang, K., et al. (2019). Downregulation of lysyl oxidase and lysyl oxidase-like protein 2 suppressed the migration and invasion of trophoblasts by activating the TGF-β/collagen pathway in preeclampsia. Exp. Mol. Med. 51, 20.

Zhou, Y., Fisher, S. J., Janatpour, M., Genbacev, O., Dejana, E., Wheelock, M. and Damsky, C. H. (1997). Human cytotrophoblasts adopt a vascular phenotype as they differentiate. A strategy for successful endovascular invasion? J. Clin. Invest. 99, 2139–2151.

