## Supplementary figures and images for "Low oxygen enhances trophoblast column growth by potentiating the extravillous lineage and promoting LOX activity"

### Supplemental Figure 1

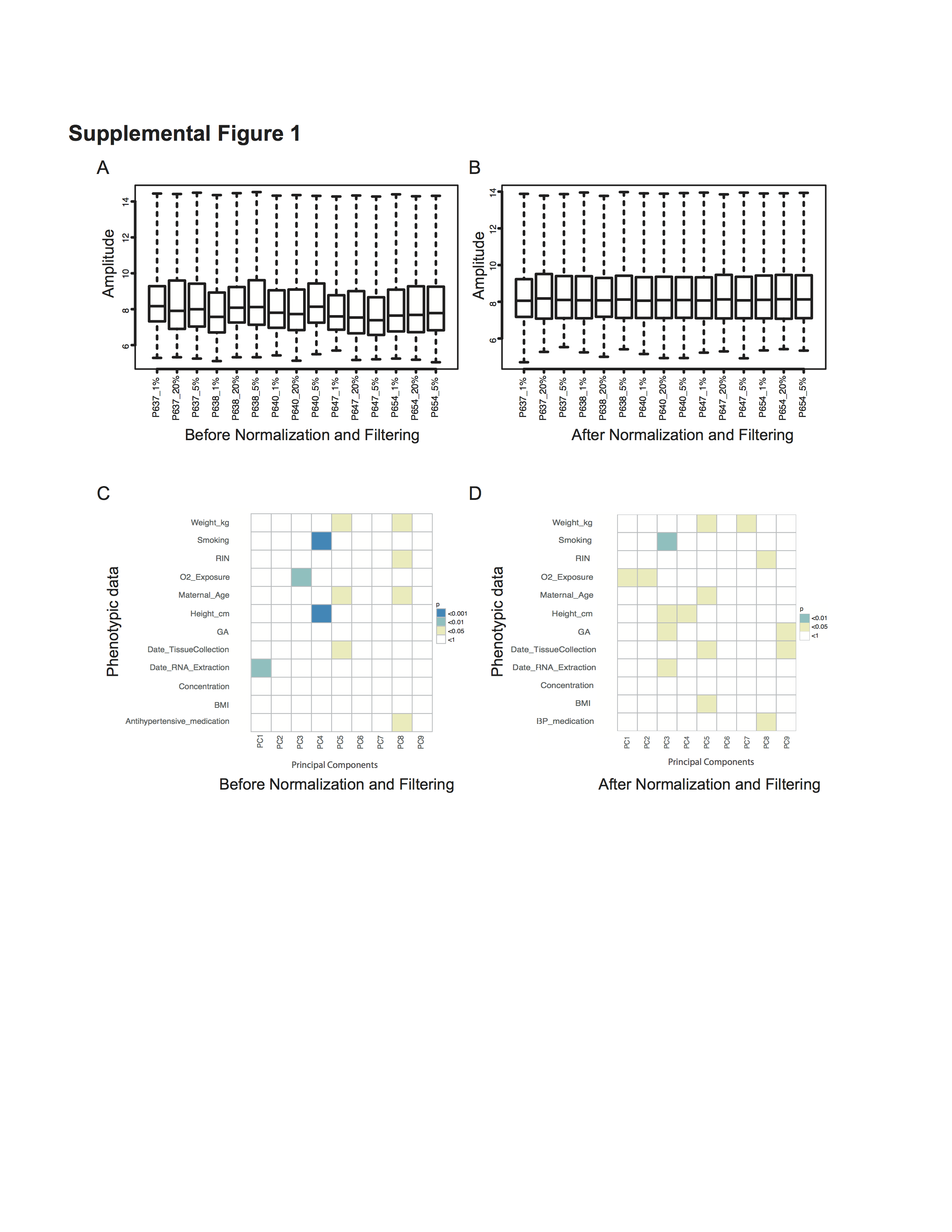

### Supplemental Figure 2

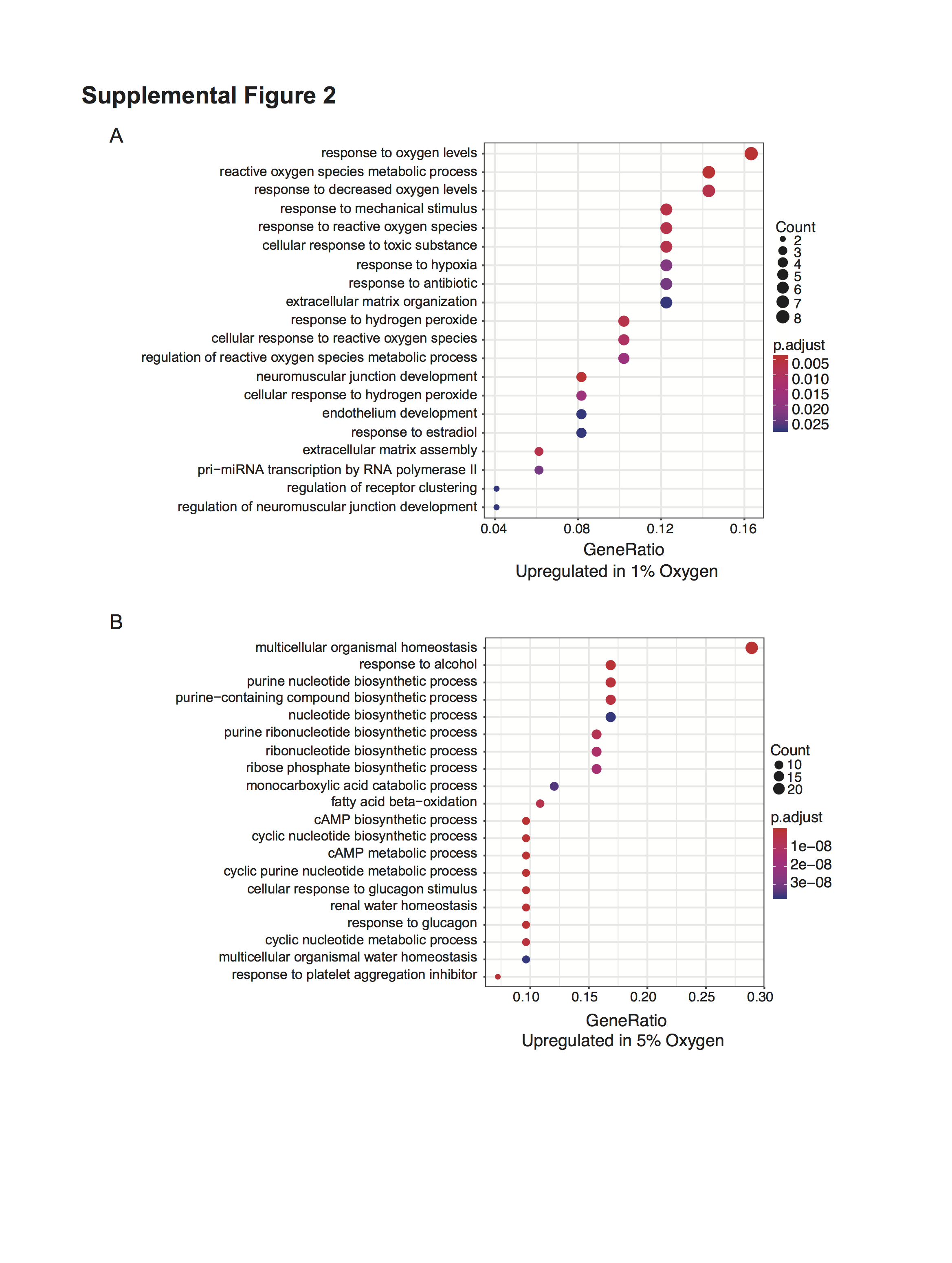

### Supplemental Figure 3

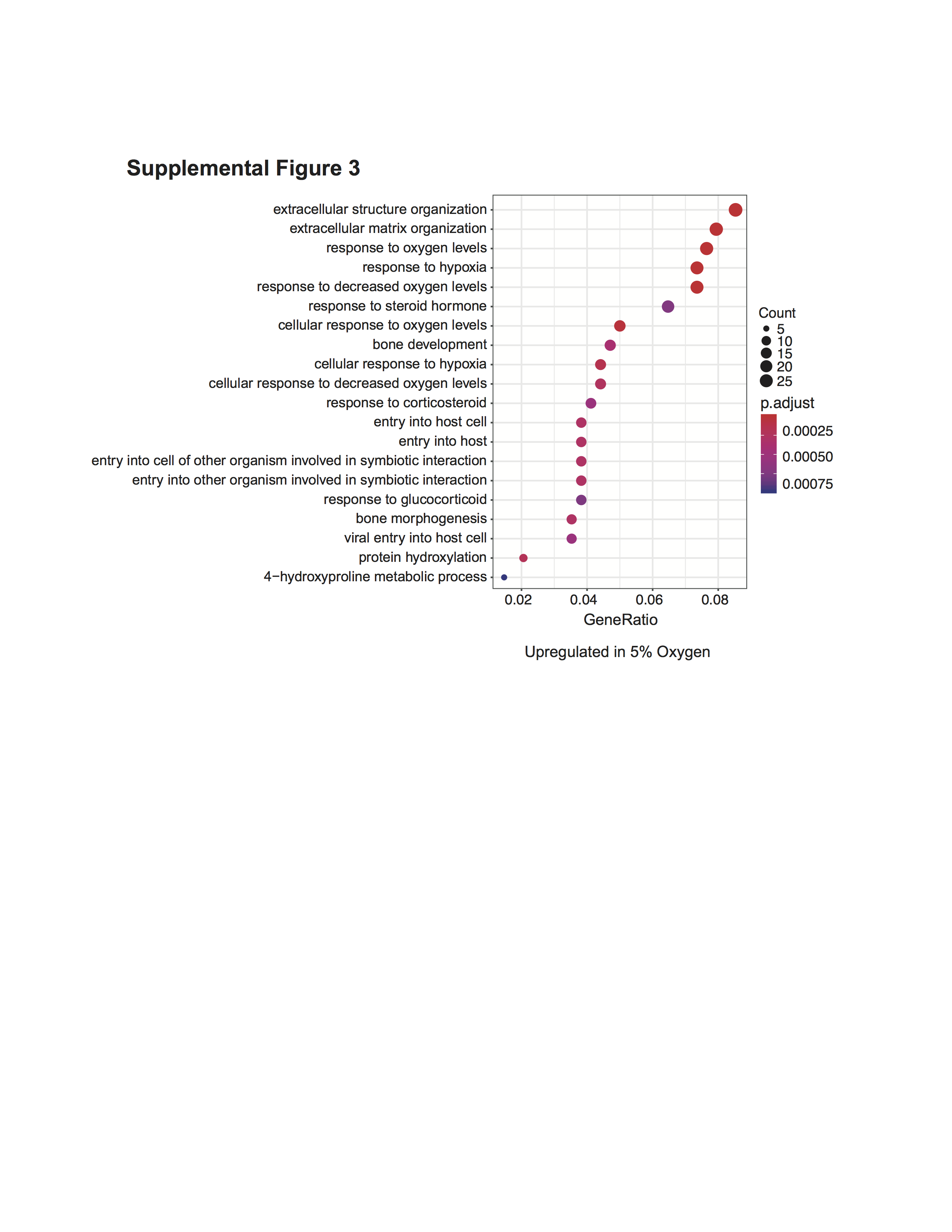

### Supplemental Figure 4

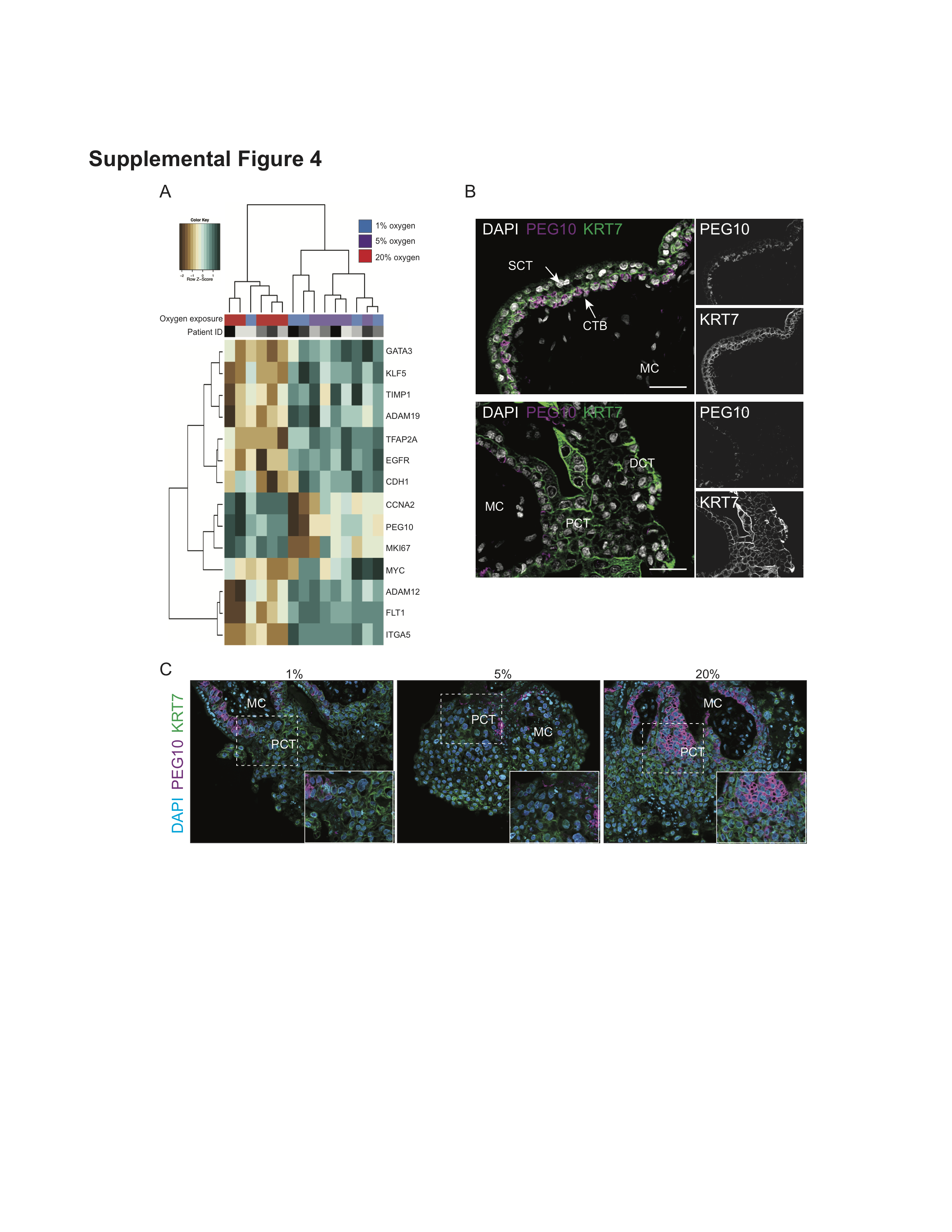
